# L-threonine mediated DAF-16/HSF-1 activation inhibits ferroptosis and increases healthspan

**DOI:** 10.1101/2022.01.24.477604

**Authors:** Juewon Kim, Yunju Jo, Donghyun Cho, Dongryeol Ryu

## Abstract

The pathways that impact longevity in the wake of dietary restriction (DR) pathway remain still ill-defined. Most studies have focused on nutrient limitation and perturbations of energy metabolism to explain the beneficial effects of DR. We showed that the essential amino acid L-threonine was elevated in *Caenorhabditis elegans* under DR, and that L-threonine supplementation increased its healthspan. To elucidate the underlying mechanism, we conducted metabolic and transcriptomic profiling using LC-MS/MS and RNA-seq analyses in worms that were fed with RNAi to induce loss of function of key candidate mediators and evaluated healthspan. L-threonine supplementation and loss-of-threonine dehydrogenase, which govern L-threonine metabolism, increased the healthspan of *C. elegans* by attenuating ferroptosis in a ferritin-dependent manner. Tran-scriptomic analysis of *C. elegans* supplemented with L-threonine showed that *FTN-1* encoding ferritin was elevated, implying *FTN-1* is an essential mediator of longevity promotion through L-threonine. Organismal ferritin levels were positively correlated with chronological aging and L-thre-onine supplementation further increased ferritin levels, which protected against age-associated ferroptosis through the DAF-16 and HSF-1 pathways. Our investigation uncovered the role of a distinct and universal metabolite, L-threonine, in DR-mediated improvement in organismal healthspan, suggesting it could be an effective intervention for preventing senescence progression and age-induced ferroptosis.

## Introduction

Reduced food intake without malnutrition can attenuate age-associated disorders in invertebrate model organisms and mammals, including humans ^1,2^. Dietary restriction (DR), defined as the coordinated reduction of intake of all dietary ingredients except vitamins and minerals, extends the lifespan of aging model organisms and improves most health phenotypes during aging ^3^. Because of diminished calorie intake during DR, most metabolic processes and metabolic intermediates are reduced, and this decreased metabolic network can induce efficient metabolic activity in many organisms ^4,5^. Thus, it is reasonable to postulate that high metabolic efficiency improves many of the physiological and molecular changes involved in age-related autonomic dysfunctions. Therefore, we speculated that increased metabolites during DR interventions might be involved in the attenuation of age-associated decline. To test this hypothesis, we screened for elevated metabolites in DR and a DR-related mutant in *Caenorhabditis elegans*. We identified several metabolic intermediates whose concentration is altered, of which the most increased was the essential amino acid L-threonine.

Nutrient amino acid imbalance has been investigated with regard of its impact on physiological processes, health and the regulation of aging. Previous studies reported that imbalanced amino acid levels may contribute to the incidence of adverse effects on aging in a rat model and suggested that reduced intake of a particular protein and amino acids may have essential roles in explaining the benefits of DR. However, the medical literature and mass media often suggest that high protein supplements, particularly those rich in essential amino acids, and having branched-chain amino acids, can overcome age-related sarcopenia, obesity, frailty, and mortality. Recent studies have been in line with this opinion, showing that specific amino acids could positively affect the organismal lifespan and health factors. For example, glycine extended the lifespan of *C. elegans* in a methionine cycle-dependent manner, and glycine supplementation extended the lifespan of male and female mice ^6,7^. Additionally, branchedchain amino acids induced a central neuroendocrine response, thereby extending healthspan ^8^, facilitating survival, and supporting cardiac and skeletal muscle mitochondrial biogenesis in middle-aged mice ^9^. These studies supported the theory that regulation of dietary amino acid levels can enhance healthspan in organisms and provided evidence in support of the further investigation of the effects of characteristic amino acids on senescence and late-life diseases ^10,11^.

In the case of L-threonine, it was reported that threonine metabolism was a crucial factor for stem cell maintenance and renewal by influencing histone methylation ^12-15^. Moreover, as a major component of mucins, threonine plays a conclusive role in the maintenance of mucosal integrity, barrier constitution, and eventually immune function regulation ^16-19^. Recently, the effects of threonine on age-associated metabolic pathways, lipid metabolism ^20^, and methylglyoxal (MGO)-mediated proteohormesis ^21^ were investigated. In this study, it was revealed that impaired threonine catabolism promoted *C. elegans* healthspan by activating a highly reactive compound in the MGO-mediated proteasome pathway. Despite these initial indications on the importance of threonine, its roles and metabolism in aging remain largely unknown.

If the effect of threonine on aging is conserved across metazoa and beyond, lifespan regulation by altered threonine metabolism should also be conserved in the commonly used aging model, *C. elegans*. We here uncovered the role of threonine and its catabolic enzyme threonine dehydrogenase (TDH) inhibition in *C. elegans* lifespan and healthspan regulation and suggest that it is mediated by ferritin-mediated ferroptosis. The use of defined diets with specific changes in amino acids composition in *C. elegans* could hence provide an experimental system with which to examine the consequences of amino acid impairment and its effects on aging and other physiological functions.

## Results

### Identification of longevity-related metabolites and their effects on *C. elegans* healthspan

To gain perspective into the longevity and change in endogenous metabolites, we sorted dietary restriction (DR)-associated metabolites for their effects on lifespan using the *C. elegans* model. Using a LC-MS/MS metabolic profiling, we detected several metabolic pathways that were significantly altered in worm subjected to the DR and DR-mimic mutant, *eat-2* model (Fig. 1a, Supplementary Data 1). According to previous studies ^1,3^, DR lowered almost all metabolic processes, because of limited calorie and nutrient intake. In contrast, we discovered that several metabolites were increased under DR and in the *eat-2* mutants compared to the wild type, N2 strain (Fig. 1b). Among the metabolites that were increased, including ribose 5-phosphate, creatinine, and glutathione, several of these metabolites were already recognized to affect longevity, including NAD^+ 22,23^ and metabolites involved lipid metabolism ^24^. Most interestingly, we found that the most elevated amino acid, threonine, delayed the aging process and extended the lifespan of nematodes by 18% (Fig. 1c, Extended Data Fig. 1a, b). The dose-response changes in mean lifespan upon threonine exposure showed that low concentrations are enhancing longevity, with 200 μM threonine exhibiting the maximal lifespan extension (Fig. 1d); 200 μM was used for subsequent assays. Threonine not only elongated lifespan, it also improved health-related parameters, average speed, and coordinated body movement (Fig. 1e), and reduced age-related triglyceride (TG) content (Fig. 1f) ^25^. Moreover, unlike many other lifespan-extending compounds, there was no significant alteration in body size, pharyngeal pumping reflected calorie intake or brood size with bacterial proliferation in the presence of threonine (Extended Data Fig. 1c-f). Importantly, threonine enhanced the levels of major reactive oxygen species (ROS) defence enzymes, superoxide dismutase (SOD), catalase activity, and resistance against acute oxidative stress with no induction of thermotolerance (Figure 1g-I, Extended Data Fig. 1g). Collectively, the DR-related metabolite threonine increased the healthspan by the induction of antioxidative protection. It was furthermore interesting to note that previous studies on anti-aging compounds and metabolites, which reported DR-mimic activity, including syringaresinol ^26^, metformin ^27^, and α-ketoglutarate ^28^, also increased the threonine content in *C. elegans* (Fig. 1j). This indicates that threonine may act as a key amino acid in the DR-mediated longevity or metabolite for healthspan extension.

**Fig 1.**
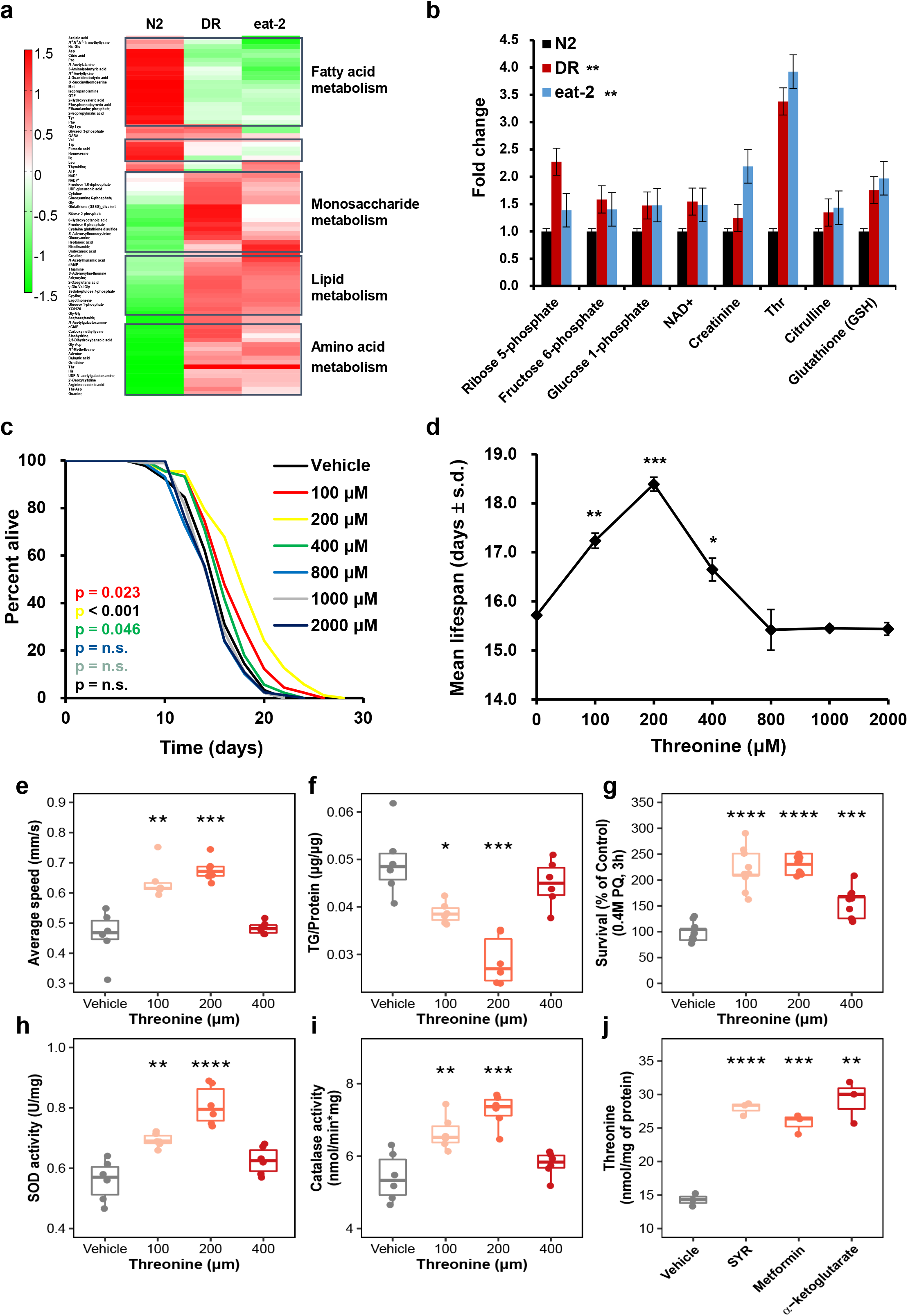
Threonine is increased in long-lived nematodes and its supplementation improves fitness and healthspan. **a**, Differentially altered metabolites, as quantified by metabolome analysis under wild-type N2 conditions, dietary restriction (DR), or the *eat-2* worms. Increased and decreased metabolites were examined by hierarchical clustering to determine the pathway and metabolites changed under the long-lived condition. **b**, Significantly elevated metabolites in DR and eat-2 compared to N2 (**p < 0.01 versus N2, two-tailed *t*-test, false discovery rate (FDR) < 0.05, n = 3 worm pellets). **c**, Effects of L-threonine (100–2000 μM) versus the vehicle (black) on lifespan (p-value listed, respectively, log-rank test); colour coding assigned to all subsequent panels. **d**, Mean lifespan of threonine treatment by concentration is depicted (*p < 0.05, **p < 0.01, and ***p< 0.001, log-rank test). **e-i**, Effects of threonine versus the vehicle in animals regarding (**e**) average speed (*p = 0.048 and **p = 0.005 versus the vehicle group, n = 10–15 worms × three assays each), (**f**) triglyceride (TG) content (*p = 0.048 and **p = 0.006 versus the vehicle group, n =3 worm pellets), (**g**) oxidative stress resistance (***p < 0.001 versus the vehicle group, n = 20 worms × nine measurements each), (**h**) superoxide dismutase (SOD) activity (*p = 0.038 and **p = 0.012 versus the vehicle group, n = 3 worm pellets), and (**i**) catalase activity (*p = 0.042 and **p = 0.008 versus the vehicle group, n = 3 worm pellets). **j**, Previously reported DR-mimic reagents also increased threonine in Day 10 worms (***p < 0.001, n = 3 worm pellets each). **e-g**, *P* values were determined by two-sided *t*-test. FDR < 0.05. Error bars represent the mean ± s.d.

### Threonine was downregulated during aging and reduced catabolism related to DR

Based on the threonine metabolic pathway (KEGG_cel00260 and wormflux_c00188), we attempted to search for age-related changes of metabolic enzymes and metabolites involving threonine (Fig. 2a). We observed an age-dependent decline of threonine content (Fig. 2b). Furthermore, DR and also *eat-2* animals exhibited decreased glycine and homoserine, which are precursors of threonine synthesis, and increased threonine without conversion (Fig. 1b, 2c). According to the Venn analysis based on public datasets, threonine metabolic enzymes, R102.4, threonine dehydrogenase (TDH)/F08F3.4, and threonine deaminase/K01C8.1 were located in the intersection of differentially expressed genes (DEG) in the DR treatment (GSE119485, padj < 0.05 in edgeR, Deseq2) ^29^ and the *eat-2* worm DEG (GSE146412, padj < 0.05 in edgeR, Deseq2) ^30^ (Fig. 2d, Supplementary Data 2). In the DR and *eat-2* mutant, threonine-producing enzyme *R102*.*4* expression was upregulated (Fig. 2e), and the expression of threonine catabolic enzyme F08F3.4 was downregulated (Fig. 2f). Compared to long-lived worms, *R102*.*4* expression was diminished in Day 5 nematodes (Fig. 2h), and *F08F3*.*4* expression increased with age (Fig. 2i). Under both conditions, the expression level of K01C8.1 did not change (Fig. 2g, j). Increased production (R102.4), together with reduced catabolism (TDH/F08F3.4) of threonine, caused an increase in threonine abundance, which potentially acted as longevity metabolites in *C. elegans*.

**Fig 2.**
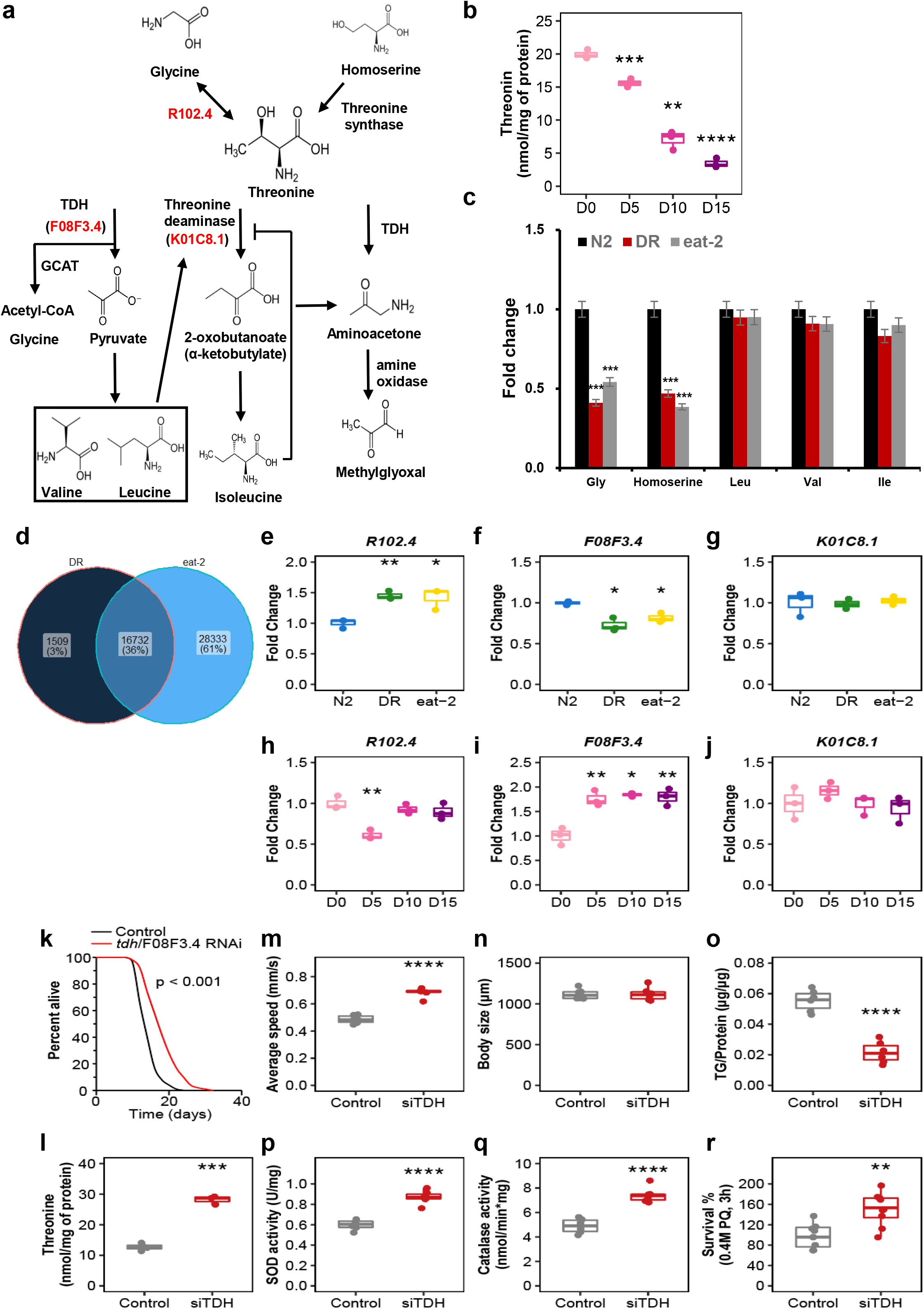
Threonine declines with age and threonine catabolism disorder enhances lifespan and healthspan. **a**, Threonine is metabolized from glycine or homoserine and catabolized to aminoacetone and methylglyoxal or a branched chain amino acid by the enzyme threonine dehydrogenase (TDH). **b**, Threonine abundance decreased with age (***p < 0.001 versus the first time point, n =3 worm pellets). **c**, Glycine and homoserine decreased, whereas leucine, valine, and isoleucine did not in DR or *eat-2* nematodes (***p < 0.001 versus N2 group, n = 3 worm pellets). **b-c**, *P* values were determined by two-sided *t*-test. FDR < 0.05. **d**, Venn analysis of DR and *eat-2* mutant using publicly available datasets. Intersection of differentially expressed genes (DEGs) of DR (p < 0.05 in edgeR, Deseq) with *eat-2* mutant DEGs compared to the wild-type N2 strain (p < 0.05). **e-g**, Among the common DEGs of DR and *eat-2* mutant, the expression of enzymes that were related with threonine metabolism were determined as (**e**) *R102*.*4* (**p < 0.01 and ***p < 0.001 versus the N2 group, n = 3 worm pellets), (**f**) *TDH/F08F3*.*4* (***p < 0.001 versus the N2 group, n = 3 worm pellets), and (**g**) *K01C8*.*1* (p > 0.05). The expression of these enzymes was measured with age as (**h**) *R102*.*4* (**p < 0.01 versus the first time point, n = 3 worm pellets), (**i**) *TDH/F08F3*.*4* (**p < 0.01 and ***p < 0.001 versus the first time point, n = 3 worm pellets), and (**j**) *K01C8*.*1* (p > 0.05). **e-j**, *P* values were determined by two-sided *t*-test. FDR < 0.05. **k**, Lifespan analysis of *tdh/F08F3*.*4* RNAi (p < 0.001, logrank test) versus control RNAi in N2 worms. **l-r**, Effects of *tdh/F08F3*.*4* RNAi versus control RNAi for (**l**) threonine contents (***p < 0.001, n = 3 worm pellets), (**m**) average speed (***p < 0.001, n = 10-15 worms × 3 assays each), (**n**) body size (p = 0.785, n = 6), (**o**) triglyceride (TG) content (***p < 0.001, n = 3 worm pellets), (**p**) superoxide dismutase (SOD) activity (**p = 0.007, n = 3 worm pellets), (**q**) catalase activity (**p = 0.002, n = 3 worm pellets), and (**r**) oxidative stress resistance (***p < 0.001, n = 20 worms × 9 measurements each). **l-r**, *P* values were determined by two-sided *t*-test. FDR < 0.05. Error bars represent the mean ± s.d.

### Threonine catabolism enzyme impairs the promotion of the healthspan of *C. elegans*

Based on increased threonine levels, we applied RNA interference (RNAi) against *tdh*/F08F3.4 (siTDH) to N2 nematodes. We attained a mean lifespan extension by 36.1% under the suppression of *tdh*/F08F3.4 expression (Fig. 2k, Extended Data Fig. 2a-c). With the concomitant increased amount of threonine (Fig. 2l), siTDH stimulated locomotion (Fig. 2m) without alteration of physical body size, pumping rate, or progeny (Fig. 2n, Extended Data Fig. 2d, e). Additionally, similar to the threonine treatment condition, siTDH lowered TG contents (Fig. 2o) and improved antioxidative defence (Fig. 2p-r) with no difference in resistance against thermal stress or bacterial growth (Extended Data Fig. 2f, g). We next determined whether blocking threonine generation affected aging parameters. The expression of the threonine-elaborating R102.4 enzyme was increased in DR with *eat-2* mutants and decreased in early-adult Day 5 worms (Fig. 2e, h). We used R102.4 RNAi to inhibit threonine production. In contrast to the observations for increased threonine (Fig. 1c-I, 2k-r), R102.4 RNAi shortened lifespan by 16.8% and reduced threonine (Extended Data Fig. 3a-e), further supporting threonine’s regulatory action in the modulation of healthspan-related factors (Extended Data Fig. 3f-o). Impairment of threonine metabolism by two different enzyme RNAi clones had adverse effects on lifespan in wild-type worms. Large amounts of threonine occurred in long-lived *C. elegans* compared to the wild-type N2 nematodes and upregulated threonine contents induced healthspan prolongation, suggesting that threonine itself enhances longevity, rather than its metabolites or precursors.

### Threonine-mediated longevity depends on transcriptional oxidative stress-response factors DAF-16 and HSF-1

To determine the molecular mechanisms of the longevity effects of threonine, we performed RNA sequencing (RNA-seq) on nematodes exposed to *tdh*/F08F3.4 RNAi for 5 d. We found 2,171 and 2,110 DEGs, respectively, which were upregulated and downregulated in comparison with control RNAi (Fig. 3a). With these remarkable changes in the transcriptome (Fig. 3b-c), impaired *tdh*/F08F3.4 expression resulted in a metabolic shift and alterations in senescence-regulatory pathways (Extended Data Fig. 4a). All DEGs were combined, and using clustered expression profiles for DEGs of DR and *eat-2* worms compared to those of the wild type nematodes, we identified 23 upregulated and 13 downregulated genes (Extended Data Fig. 4b, c). Focusing on the transcription factors of these DEGs and given the distinguished role of lifespan, we next tested whether *tdh*/F08F3.4 has an epistatic interaction with DR, insulin/IGF-1, or stress-response regulators. Similar to the results for increased threonine and decreased *tdh*/F08F3.4 levels in DR and *eat-2* mutant (Fig. 1b, 2f), siTDH and threonine application failed to further extend a lifespan of *eat-2*(*ad465*), which was different from N2 nematodes (Fig. 3d, Extended Data Fig. 4d-e, 5a-c). Interestingly, threonine catabolism impairment, or threonine itself, interacted with *daf-16*, although its upstream transcription factor *daf-2* did not (Fig. 3e-f, Extended Data Fig. 4f-i). This requirement of *daf-16* was specific because the lifespan of the even longer-lived insulin/IGF-1 receptor *daf-2* mutant worms was further prolonged by siTDH and threonine (Extended Data Fig. 5d-i). Lastly, and according to a previous study with another threonine catabolic enzyme, glycine-C-acetyltransferase (GCAT) ^21^, transcriptional stress-response regulator HSF-1 was also considered. The higher antioxidative enzyme activity and oxidative stress resistance in threonine-treated animals suggested that threonine-mediated longevity may involve a similar context to that initiated by stress stimulation. Furthermore, we confirmed that lifespan extension effects of both siTDH and threonine intervention were fully negated in *hsf-1*(*sy441*) worms, which lacked the transactivation domain of the protein (Fig. 3g, Extended Data Fig. 4j-k, 5j-l). These findings demonstrated epistatic interplay of threonine with *daf-16* and *hsf-1*. In addition we confirmed that the nuclear translocation and increased expression of *daf- 16::gfp* or *hsf-1::gfp* was intensely stimulated by *tdh*/F08F3.4 RNAi and additional threonine (Fig. 3h-k).

**Fig 3.**
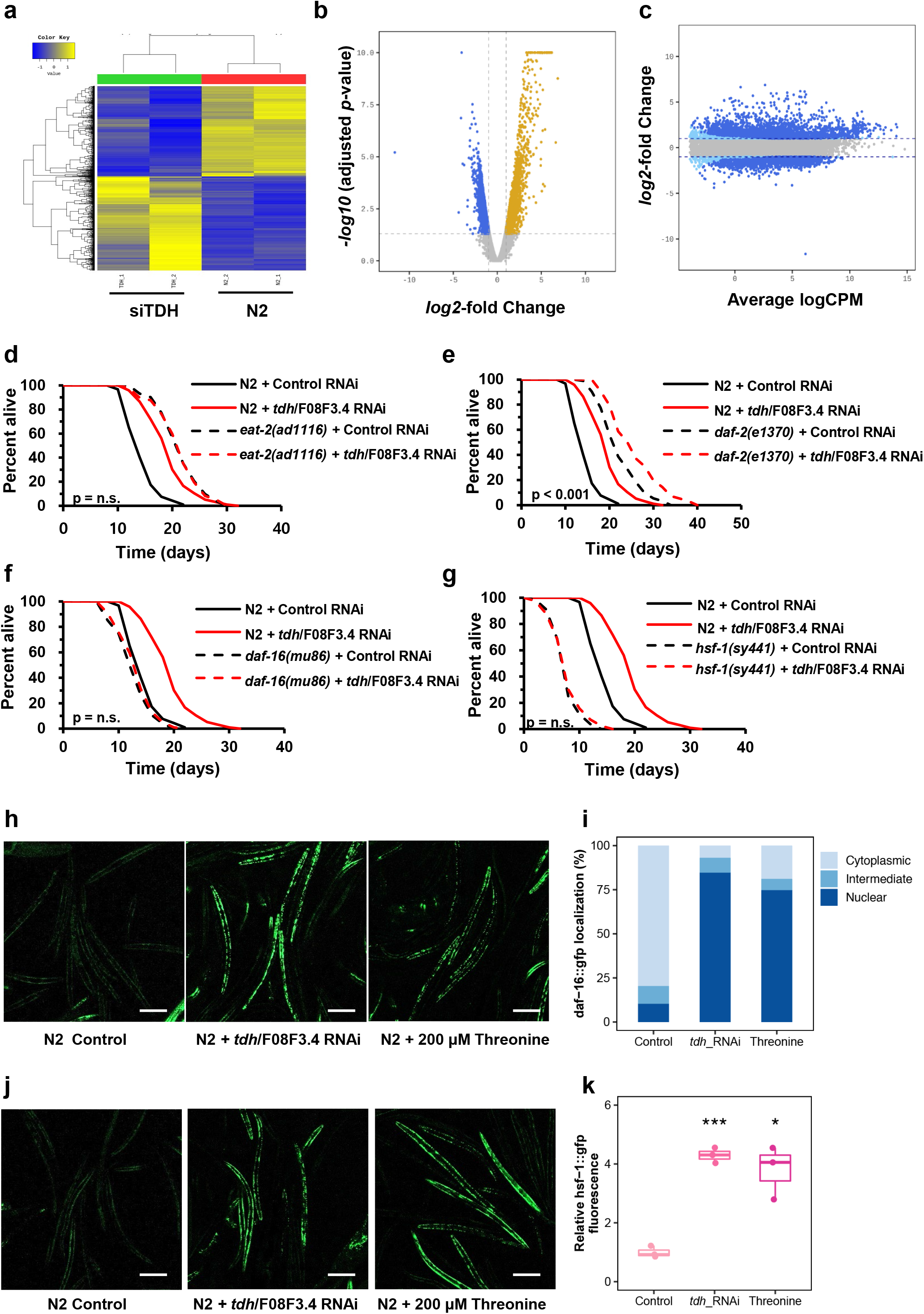
Lengthening of health span by *tdh/F08F3*.*4* RNAi and threonine supplement is mediated by transcription factors DAF-16 and HSF-1-dependent metabolic changes. **a**, Gene clusters whose expression is regulated by *tdh/F08F3*.*4* RNAi versus the wild-type N2 are represented by hierarchical clustering analysis (Euclidean distance, complete linkage). **b**, DEGs in *tdh/F08F3*.*4* RNAi compared to the N2 visualized with a volcano plot. An adjusted p value < 0.05 and ❘log_2_ (*tdh/F08F3*.*4* RNAi / N2 ratio)❘ > 2 were applied to define significantly up-or downregulated genes (blue and yellow dots, respectively) or genes without change in expression (grey dots). **c**, DEGs as quantified by transcript levels by deep sequencing analysis under treatment with *tdh/F08F3*.*4* RNAi; grey dots represent no differential regulation; dark blue and light blue dots depict DEGs according to edgeR analysis (dark blue: ❘FC❘ ≥ 2 with p < 0.05; light blue ❘FC❘ ≥ 2). **d-g**, Lifespan assay of *tdh/F08F3*.*4* RNAi (red-dashed) versus control RNAi (black-dashed) for (**d**) *eat-2*(*ad465*) (p = 0.786), (**e**) *daf-2*(*e1370*) (p < 0.001), (**f**) *daf-16*(*m26*) (p = 0.627), and (**g**) *hsf-1*(*sy441*) (p = 0.492) mutant worms with survival curves of N2 (Bold-line). **d-g**, *P* value determined by two-sided log-rank test. **h**, Expression and localization (cytoplasm/intermediate/nuclear) of DAF-16::GFP under control conditions, after exposure to *tdh/F08F3*.*4* RNAi or 200 μM threonine treatment for 5 d, and **i**, localization quantifications (***p < 0.001, *χ*^2^ test). **j**, Expression of HSF-1::GFP at control, *tdh/F08F3*.*4* RNAi, or 200 μM threonine, and **k**, corresponding quantifications (***p < 0.001, *χ*^2^ test). Scale bars, 200 μm. Error bars represent the mean ± s.d.

### FTN-1 expression is required for threonine-mediated longevity

Based on these results, we selected six and two commonly up- and downregulated genes, respectively, which are transcribed by the transcription factor DAF-16 or HSF-1, from the intersecting genes of *tdh*/F08F3.4 RNAi, DR, and *eat-2* worms (Supplementary Data 3). All of these genes were individually targeted by respective RNAi for worms with the exception of genes already examined for threonine metabolism, R102.4 and *tdh*/F08F3.4. We investigated lifespan changes by applying RNAi of these four genes: glutathione *S*-transferase, *gst-19* (Fig. 4a, Extended Data Fig. 6a-b); inferred organismal homeostasis of metabolism variant, T12D8.5 (Fig. 4b, Extended Data Fig. 6c-d); *Caenorhabditis* bacteriocin, *cnc- 2* (Fig. 4c, Extended Data Fig. 6e-f), and ferritin, *ftn-1* with *tdh*/F08F3.4. The RNAi revealed that only *ftn-1*/C54F6.14 exhibited an interaction with the impaired *tdh*/F08F3.4 expression (Fig. 4d, Extended Data Fig. 6g-h). *tdh*/F08F3.4 RNAi and threonine were furthermore unable to additively extend the lifespan under the *ftn-1* null mutant, *ftn-1*(*ok3625*) (Fig. 4e-f, Extended Data Fig. 6i-l), indicating that the conserved iron-cage protein, ferritin 1 (FTN-1) is essential for threonine-mediated longevity.

To determine the significance of FTN-1 to threonine-related longevity, we measured *ftn-1* expression with physiological aging and in N2 worms fed with *tdh*/F08F3.4 RNAi. As previously reported ^31^, *ftn-1* expression increased with age, but more importantly, its expression was further elevated by the reduced expression of *tdh*/F08F3.4, especially at early adult stages (Fig. 4g-h). Thus, FTN-1 is required for the longevity advantage of threonine. Given the interaction of *ftn-1* expression with several genes, we exposed DR and *eat-2* worms to the target gene RNAi, with control RNAi serving as control. Although von Hipple-Lindau protein 1 (*vhl-1*), hypoxia-inducible transcription factor (*hif-1*), and prolyl hydroxylase *egl- 9* had no effect on *ftn-1* expression, *daf-16* and *hsf-1* did affect elevated *ftn-1* expression in DR and *eat-2* worms, compared with the wild-type nematodes (Extended Data Fig. 7a, b). This may have occurred because of the specific regulation of *ftn-1* in the threonine-mediated longevity condition. Because *ftn-1* expression is modulated by oxygen concentration, *ftn-1* was regulated differently under normoxia, hypoxia, and hyperoxia conditions using specific mutations of the target genes (Extended Data Fig. 7c-e). Similar to the DR and *eat-2* animals, *daf-16* and *hsf-1* were necessary to induce *ftn-1* expression by *tdh*/F08F3.4 RNAi, indicating threonine-induced longevity is impacted by increasing organismal levels of FTN-1.

**Fig 4.**
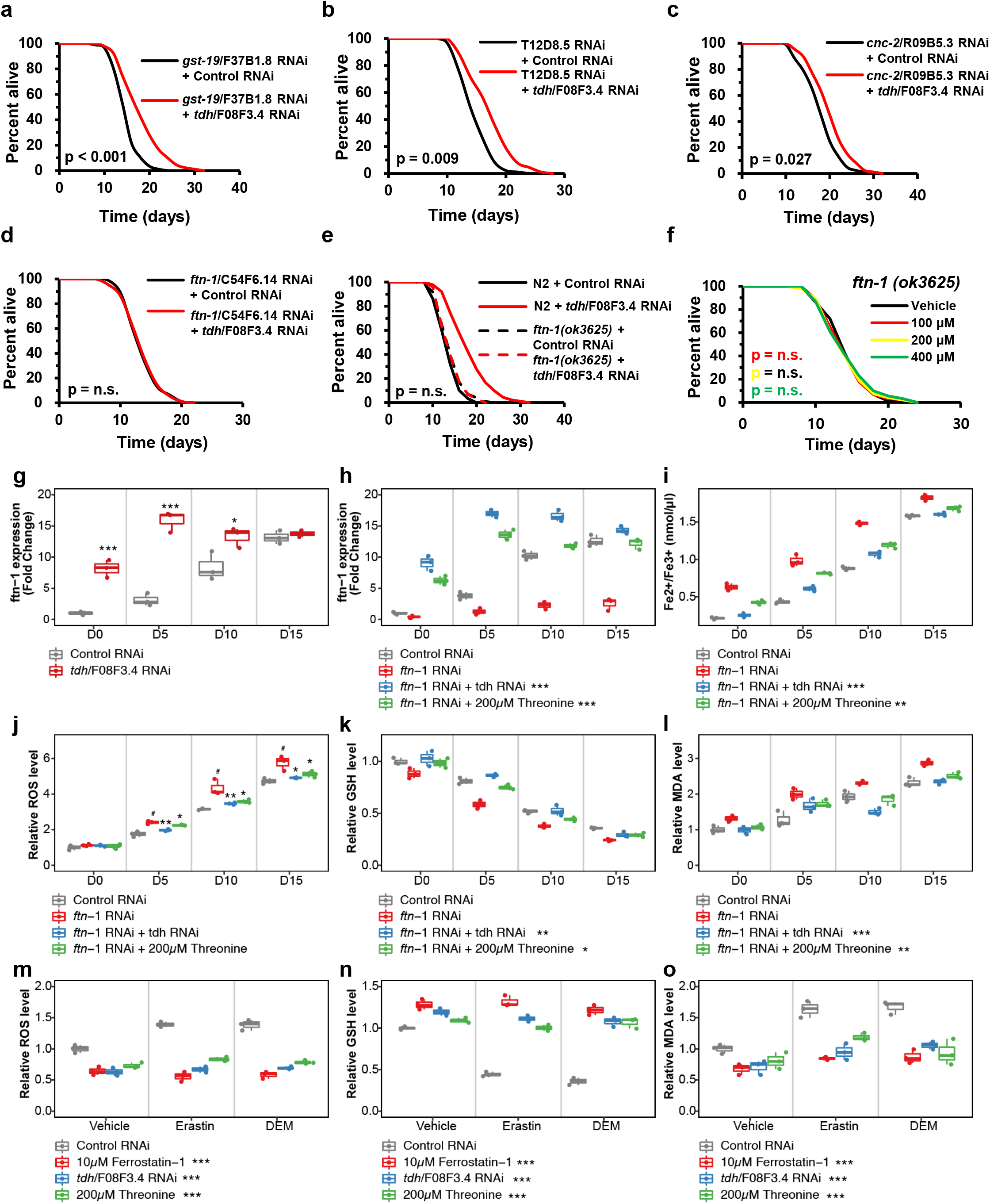
Increasing threonine manipulations upregulate expression of ferritin protein, FTN-1, and FTN-1-mediated ferroptosis suppression. **a-d**, Lifespan assay of *tdh/F08F3*.*4* RNAi (red) versus control RNAi (black) with transcripts of DAF-16 or HSF-1 among the commonly regulated genes of *tdh/F08F3*.*4* RNAi, DR, and *eat-2* (by Venn analysis, Supplementary Data 3), (**a**) *gst-19/F37B1*.*8* RNAi, (**b**) *T12D8*.*5* RNAi, (**c**) *cnc- 2/R09B5*.*3* RNAi, and (**d**) *ftn-1/C54F6*.*14* RNAi worms. **e**, Lifespan assay of *ftn-1(ok3625)* with *tdh/F08F3*.*4* RNAi (red-dashed) versus control RNAi (black-dashed) compared with survival curves of N2 (bold-line). **f**, Effects of threonine (100, 200, and 400 μM) versus the vehicle (black) on lifespan of the *ftn-1(ok3625)* mutant. *P* value and lifespan assay data summarized in Extended Data Table 1. **g**, Expression of *ftn-1* under *tdh/F08F3*.*4* RNAi was determined at intervals across lifespan (*p < 0.05 and ***p < 0.001 versus control RNAi, n = 3 worm pellets). **h**, Expression levels of *ftn-1* under *ftn-1*/C54F6.14 RNAi (#p < 0.001 compared to control RNAi, ***p < 0.001 versus *ftn-1* RNAi, n = 3 worm pellets). **i**, Quantification of Fe^2+^ /Fe^3+^ iron contents of nematode with *ftn-1*/C54F6.14 RNAi at intervals across lifespan (#p < 0.001 compared to control RNAi, **p < 0.01 and ***p < 0.001 versus the *ftn-1* RNAi, n = 3 worm pellets). **j**, Relative Amplex Red fluorescence in supernatant of worms (#p < 0.001 compared to control RNAi, *p < 0.05 and **p < 0.01 versus *ftn-1* RNAi, n = 3 worm pellets). **k**, Total glutathione (GSH) level was normalized to the GSH level in worms exposed to *tdh/F08F3*.*4* RNAi or threonine (#p < 0.001 compared to control RNAi, *p < 0.05 and **p < 0.01 versus *ftn-1* RNAi, n = 3 worm pellets). **l**, Levels of the lipid peroxidation end product, malondialdehyde (MDA), were measured and normalized against the mean of control RNAi-treated worms for independent samples (#p < 0.001 compared to control RNAi, **p < 0.01 and ***p < 0.001 versus the *ftn-1* RNAi, n = 3 worm pellets). **m-o**, Effects of *tdh/F08F3*.*4* RNAi or threonine under the ferroptosis inducer erastin or diethyl maleate (DEM) by GSH depletion as relative levels of (**m**) ROS, (**n**) GSH, and (**o**) MDA with ferroptosis inhibitor, ferrostatin-1 (***p < 0.001 versus control RNAi, n = 3 worm pellets). Overall differences between conditions were analysed by two-way ANOVA. Differences in individual values or between two groups were determined using two-tailed *t*-tests (95% confidence interval). Error bars represent the mean ± s.d.

### FTN-1 mediates threonine-dependent ferroptosis alleviation

We next estimated whether iron contents might contribute to the long-lived phenotype. Similar to humans, *C. elegans* FTN-1 is induced by iron, and accumulated reactive ferrous iron results in a non-apoptotic form of cell death, ferroptosis ^32^. Quantifying the rate of ferrous iron (Fe^2+^, redox-active) / ferric iron (Fe^3+^, stable) concentrations following *tdh*/F08F3.4 RNAi or threonine also indicated a global diminution in organismal age-dependent ferroptosis (Fig. 4i, Extended Data Fig. 7f). Decreased levels of ferrous iron may result from expended iron retention of FTN-1 and result in inactivating ferroptosis. Moreover, this redox-active iron ratio (Fe^2+^/ Fe^3+^) was also dependent on *daf-16* and *hsf-1*, similar to the regulation of *ftn-1* expression in DR or *eat-2* nematodes (Extended Data Fig. 7g, h). In the ferroptosis pathway, ferrous iron accumulates over the adult lifetime of *C. elegans* ^33^ and exists as abundant reactive ferrous iron, which could generate massive damage through ROS, eventually inducing ferroptosis and shortening the lifespan ^34,35^. Hence, with the changes accruing during physiological aging, we detected in worms fed with *tdh*/F08F3.4 RNAi or with reduced threonine an associated increase in ROS levels in whole nematode lysates, particularly during the late adult stage (Fig. 4j, Extended Data Fig. 7i). Additionally, we confirmed that the absence of *daf-16* or *hsf-1* failed to reduce ROS levels in DR and *eat-2* worms, analogous to *ftn-1* expression (Extended Data Fig. 8a-c). As a substrate for the main regulator of ferroptosis glutathione peroxidase 4 (GPX4), glutathione (GSH) deficiency is coupled to ferrous iron accumulation in nematodes, and both occur to irritate cells leading to ferroptosis ^33^. To systematically examine this, we determined whole GSH levels in worm lysates after *tdh*/F08F3.4 RNAi or threonine feeding at time points during aging. With the age-response decline in GSH levels, threonine increasing interventions prevented GSH depletion (Fig. 4k, Extended Data Fig. 7j) and increased the GSH level in DR and *eat-2* worms, which was modulated by *daf-16* or *hsf-1* as assayed by the target gene RNAi (Extended Data Fig. 8d-f). Redox-active iron-dependent lipoxygenases and iron-catalysed peroxyl radical-mediated autoxidation could induce lipid peroxidation, which executed ferroptosis ^36^. We examined the toxic lipid peroxidation end-product malondialdehyde (MDA) to assay the ferroptosis level ^37^, *tdh*/F08F3.4 RNAi or threonine feeding reduced MDA levels in an age-dependent manner (Fig. 4l, Extended Data Fig. 7k). Similarly, the MDA level was lowered dependent on *daf-16* or *hsf-1* in DR and *eat-2* nematodes (Extended Data Fig. 8g-i).

This, together with the results summarized above, raised the hypothesis that increasing threonine interventions attenuate compromised ferritin capacity, inhibiting redox-active iron-mediated ferroptosis. To test this hypothesis, we determined the inhibitory effects of threonine elevation treatment against ferroptosis stimulated by introducing *ftn-1*/C54F6.14 RNAi. As expected, with recovery of *ftn-1* expressions (Fig. 4h), increasing threonine application reduced ferroptosis markers under *ftn-1* depletion status. Treatment of *tdh*/F08F3.4 RNAi or threonine supplementation lessened redox-active iron ratio, ROS levels, GSH levels, and also MDA levels in whole nematode lysates in FTN-1-dependent manner (Fig. 4i- l). Furthermore, we confirmed the suppressive effects of increasing threonine treatment against ferroptosis stimulated by using acute depletion of GSH by diethyl maleate (DEM) ^38^ or a glutamate/cystine antiporter suppressor, erastin ^39^. The results provided convincing evidence for the anti-ferroptosis activity of *tdh*/F08F3.4 RNAi or threonine supplementation against exposure to ferroptosis-inducing agents as reflected by reduced ROS, increased GSH, and reduced MDA levels (Fig. 4m-o). Additionally, we assessed the age-associated pigment, lipofuscin, which is defined as iron-filled intracellular inclusions triggering ferroptosis ^40^. Lipofuscin accumulation diminished with threonine or *tdh*/F08F3.4 RNAi feeding but enhanced with R102.4 RNAi (Extended Data Fig. 8j, k). When restoring the cellular GSH level by threonine interventions, we measured the highly reactive by-product of glycolysis, MGO, with its scavenger, GSH. MGO is one of the representative cell-permeant precursors of advanced glycation end-products (AGEs), which are associated with several age-related diseases. The MGO/GSH ratio was lowered with manipulations that increased threonine levels; indicating, it was beneficial for longevity (Extended Data Fig. 8l-n). Furthermore, the MGO/GSH ratio increased with aging and long-lived DR and *eat-2* worms showed a greatly decreased MGO/GSH ratio and a concordantly increased lifespan (Extended Data Fig. 8o-p).

In the current study, we focused on threonine as a commonly increased metabolite in DR-mediated long-lived worms. Blocking threonine dehydrogenase or supplementation with threonine itself lead to increased organismal threonine content. Increased threonine content leads through a DAF-16 and HSF-1 dependent mechanism to the activation of FTN-1 and the antioxidant system, which attenuates ferroptosis, and mediates lifespan extension and healthspan-contributing factors in *C. elegans* (Fig. 5).

**Fig 5.**
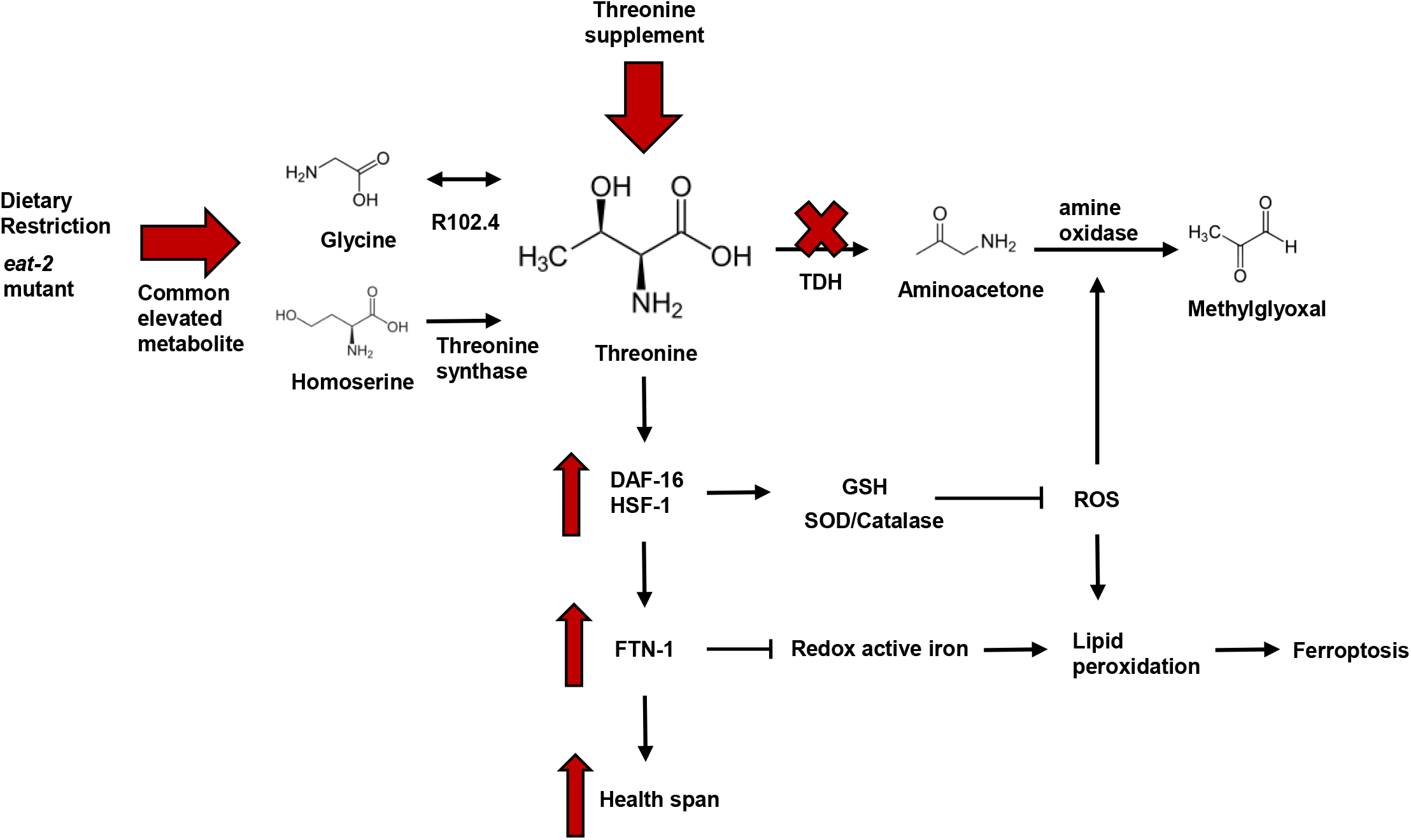
Mechanistic model of threonine-mediated longevity. Summary of the effects of *tdh/F08F3*.*4* RNAi or threonine supplementation on DAF-16/HSF-1 and ferroptosis signalling results in healthspan extension. Threonine catabolism impairment by *tdh/F08F3*.*4* RNAi leads to increased threonine and decreased formation of methylglyoxal. Threonine increased SOD/catalase activity and GSH expression by DAF-16/HSF-1 activation. Subsequently, and dependent on DAF-16 and HSF-1, activation of ferritin protein, FTN-1, reduced redox active iron contents and eventually ferroptosis, which mediates lifespan extension and health parameters.

## Discussion

We specified commonly increased metabolites that may affect longevity in DR-mediated long-lived animals through a metabolomics approach. Among the macronutrients, we identified threonine as most increased amino acid, despite previous studies showing that limitation of proteins or amino acids is the most forceful single molecular restriction beneficially impacting on longevity and age-associated diseases ^8,41^. Besides its role in protein metabolism, threonine is also impacting on organismal maintenance processes, such as the synthesis of immune proteins and production of intestinal mucus layers ^19,42,43^. According to whole-genome sequencing analysis of semi-supercentenarians in Italy, glycine, serine, and threonine metabolism were identified as the top-tier pathways involved in longevity ^44^. Furthermore, a recent multi-omics study in mice revealed that the threonine pathway was involved in longevity and related molecular mechanisms ^45^. Together these studies indicated that threonine or its metabolites may affect organismal health- and life-span.

Impairing threonine catabolism has been reported to promote the healthspan of *C. elegans*. This study concentrated on investigating the action of glycine C-acetyltransferase (GCAT), which is involved in the conversion of the intermediate 2-amino-3-ketobutyrate to glycine and its impairment resulting in an increment in MGO-mediated proteohormesis ^21^. Here, we focused on threonine contents itself. We investigated the role of threonine and enzymes involved in its metabolism, which directly influence threonine abundance. First, we determined whether threonine could affect *C. elegans* lifespan or it was only a consequence of longevity. Low-dose threonine enhanced lifespan and healthspan-related parameters with improved antioxidative enzyme activities. Because DR-mediated longevity has been correlated with reduced protein and amino acid intake, these effects raise an intriguing question about how certain amino acids have the potential to promote healthspan. Although many of studies have reported the benefits of amino acid restriction, recent research has provided clues about the longevity advantages of specific amino acids in many organisms. Supplementation of the single amino acid glycine extended lifespan of *C. elegans* ^7^, *Drosophila* ^46^, and mice ^6^ by attenuation of the progression of dystrophic pathologies. In addition, branched-chain amino acids, a group of essential amino acids, such as leucine, isoleucine, and valine, could impact health and lifespan by affecting amino acid balance ^8,9,47^. Incited by these findings, we investigated the merits of threonine on fitness and lifespan. Based on published RNA-seq data, the threonine synthesis enzyme, R102.4, was shown to be upregulated in DR-related mutant *eat- 2*(*ad465*) worms and TDH/F08F3.4 was identified amongst the DEGs in the long-lived mutants, *daf-2*(*e1370*) and *eat-2*(*ad465*) ^7,48^. Both threonine synthesis using glycine and homoserine and inhibition of the threonine catabolic enzyme TDH increase threonine contents in DR-mediated longlived worms.

We showed that threonine upregulates ferritin, FTN-1, expression. Although the total amount of ferritin protein is increased, the iron storage capacity of ferritin is compromised with age and ablated with senescence ^33,49^. We assumed that threonine increases healthspan by attenuating agerelated ferroptosis. Increasing threonine levels upregulates FTN-1 renewal through a mechanism involving the transcription factors DAF-16 and HSF-1, subsequently reduces the hallmarks of ferroptosis, the redox-active iron level. and blocks the generation of. Moreover, we detected ferroptosis-related factors, age-pigment lipofuscin accumulation and highly reactive aminoacetone, a precursor of MGO, found to induce dose-dependent ferrous iron release from ferritin ^50^ decreased in threonine-associated longevity.

According to chromatin immunoprecipitation sequencing (ChIP-seq) results, it has been reported that human ferritin, ferritin heavy chain 1 (FTH- 1), expression is initiated by the activation of the human homolog of DAF-16, FoxO3 ^51^. This indicated that ferritin expression might also be regulated by threonine modulation in humans through the activation of the FoxO3 transcription factor. Ferritin imbalance provokes diverse health disorders and diseases, such as iron-deficiency anaemia, hypothyroidism, celiac disease in case of deficiency, adult-onset Still’s diseases, porphyria, and an acute inflammatory reaction in some cases of COVID-19 ^52 53^. These observations and manifestations associated with ferritin homeostasis and its turnover are crucial factors to human health and organismal energy balance ^54^. So far most of the positive effects of threonine are linked to enhanced immunity, through mucus formation ^18,42^, and stimulation of pluripotency of embryonic stem cells ^12-15^. Here, we propose another angle how threonine can achieve health- and lifespan benefits in *C. elegans* through the attenuation of cellular ferroptosis. Our work should further additional studies to evaluate whether these effects on fitness and longevity translate to mammals and humans.

## Online content

Any method, additional references, extended data, supplemental information can be found with this article online.

## Acknowledgements

*C. elegans* strains used in this research were provided by the *Caenorhabditis* Genetics Center (University of Minnesota, USA), which is funded by the NIH Office of Research Infrastructure Programs.

## Author Contributions

J.K. created the experimental design and performed all experiments, except as stated otherwise. Y.J. contributed to analysis of metabolomics and RNA sequencing data. D.C. performed experiments of isolated nematodes. J.K. and D.R. designed the study and wrote the manuscript; D.R. edited the manuscript; and J. K. and D.R. conceived and designed the research, analysed and interpreted the data, and administered the project. All authors discussed the results and commented on the manuscript.

## Additional information

**Supplementary information** The online version contains supplementary material available

**Correspondence and requests for materials** should be addressed to J.K and D.R.

## Methods

### Nematode strains and maintenance

*C. elegans* strains were maintained on Nematode Growth Media (NGM) agar plates at 20°C using OP50 bacteria as a food source. The following strains were used (strain, genotype): Bristol N2, wild type; DA1116, *eat-2(ad1116)*II; CB1370, *daf-2(e1370)*III; CF1038, *daf-16(mu86)*I; PS3551, *hsf-1(sy441)*I; TJ356, zIs356 [daf-16p::daf-16a/b::GFP+rol-6(su1006)]; OG532, *<u>hsf-1</u>*(*<u>sy441</u>*)I; <u>drSi13</u>II [hsf-1p::hsf-1::GFP::unc-54 3’UTR+Cbrunc-119(+)]; RB2603, *ftn-1*(*ok3625*)V. They were all obtained from the Caenorhabditis Genetics Center (CGC). All strains are listed in Extended Data Table 1.

### Worm Synchronization and food preparation

To obtain synchronized nematode, six unmated hermaphrodites were placed on an NGM plate with sufficient food. After 4 days, lysed F1 progenies that were selfing hermaphrodites yielded synchronized hermaphrodite eggs. The obtained eggs were grown on NGM plates. OP50 bacteria were cultured at 37 °C in LB medium for up to 12 hours. The OP50 concentration was calculated by a combination of counting the colonies and measuring the OD at a wavelength of 600 nm. The OP50 was centrifuged, resuspended and diluted in H_2_O at bacteria concentrations of 10×10^11^ bacteria ml^-1^ or 10×10^9^ for the dietary restriction (DR) experiments until day 4 of feeding beginning at day 1 of adulthood, same condition with previous study which of RNAseq data used in this study (GSE119485).

### Metabolites extraction and metabonomic profile of *C. elegans*

LC-MS/MS-based metabonomic profile of frozen worm pellets was performed (Human Metabolome Technologies, Yamagata, Japan). The wild type N2, DR treatment, or *eat-2* mutant of one-day-old animals were raised over a period of 4 days. Frozen worm samples were mixed with 800 μl of 50% acetonitrile (v/v) containing internal standards (20 μM for cation measurement and 5 μM for anion measurement), homogenized in the presence of beads, and centrifuged. The supernatant was then filtered through 5-kDa cut-off filters (Ultrafree-MC PLHCC, Merck KGaA, Darmstadt, Germany) to remove macromolecules. The filtrate was centrifugally concentrated and diluted in 25 μl of ultrapure water for CE/TOF-MS analysis. For LC/TOF-MS analysis, frozen worm samples were mixed with 500 μl of 1% formic acid in acetonitrile (v/v) containing internal standards, homogenized, and centrifuged. The supernatant was filtered through a 3-kDa cut-off filter (Nanocep 3K Omega, Pall Corporation, MI, USA) to remove proteins and then passed over a column (Hybrid SPE phospholipid 55261-U, Supelco, PA, USA) to remove phospholipids. The column eluate was desiccated and suspended in 100 μl of 50% isopropanol. Three independently collected worm pellets were used for each assay with CE-TOF/MS system and 1200 series Rapid Resolution system SL, subsequently LC/MSD-TOF (Agilent Technologies, CA, USA).

### Assay for threonine levels in *C*.*elegans*

Synchronized adult worms were collected, washed 3 times with M9 buffer, and flash grounded frozen pellets. Worms were lysed with lysing matrix C tubes (#6912-100, MP Biomedicals, CA, USA) and the TissueLyser II (Qiagen, Hilden, Germany) high speed bench-top homogenizer in the 4°C room with condition of 30 Hz for 2 min, then rest on ice for 1 min, repeated 3 times. Lysed worms were centrifuged (13,200 rpm, 10 min, 4°C) to remove debris, and supernatant was used. Threonine contents were measured with threonine assay kit (ab239726, Abcam, MA, USA) with normalization with total protein concentration (BCA assay, Pierce, IL, USA).

### Preparation of threonine supplementary plates

The amino acid threonine was acquired (Sigma-Aldrich, MO, USA) and stock of 1 M was made by dissolving in water. The pH of stock solution was adjusted to 6.0 - 6.5, the same pH as the vehicle plates, with sodium hydroxide and threonine stock solution sterilized with 0.45 μm Millipore filter (Millipore, MA, USA) before use. Threonine treatment was performed by supplementing the compound in heat-inactivated OP50 (65°C, 30 min with 1M magnesium sulfate and 5 mg/ml cholesterol), unless it applied with RNAi clones.

### RNAi in *C. elegans*

For RNAi-mediated gene knockdown, we fed worms HT115 (DE3) expressing target gene double stranded RNA (dsRNA) from the pL4440 vector. The clones for RNAi against genes were obtained from the *C. elegans* ORF-RNAi Library (Thermo Scientific, Horizon Discovery, MA, USA). All clones were verified by sequencing (Macrogen, Seoul, Korea) and efficient knockdown was confirmed by quantitative RT-PCR (qRT-PCR) of the mRNA. The primer sequences used for qRT-PCR are provided in Extended Data Table 2. Bacteria were spotted on NGM plates containing additionally 1 mM isoprophyl-β-D-thiogalactoside (IPTG) (Sigma-Alrich, MO, usa). Incubation with dsRNA initiated 64 hours after population synchronization and then young adult worms transferring to the respective treatment plates. We confirmed the RNAi knockdown of *tdh* by qRT-PCR and transcripts levels of *tdh* was reduced by 71% in larvae that were cultivated on corresponding dsRNA-expressed bacteria.

### Lifespan analysis

Lifespan assays were carried out at 20°C on NGM agar plates using standard protocols and were replicated in three independent experiments. *C*.*elegans* were synchronized with 70 μl M9 buffer, 25 μl bleach (10% sodium hypochlorite solution) and 5 μl of 10N NaOH. Either a timed 64 hours of egg lay or egg preparation, around 150 young adults were transferred to fresh NGM plates containing 49.5 μM 5-fluoro-2’-deoxyuridine (FUDR, Sigma-Aldrich, MO, USA) to prevent progeny production. All compounds were blended into the NGM media after autoclaving and before solidification. Living worms were transferred to assay plates, which were seeded with OP50 or a designated RNAi feeding clone with 50 μg ml^-1^ ampicillin (Sigma-Aldrich, MO, USA) every other day. Worms that crawled of the plates or exhibited internal progeny hatching were censored and showed no reaction to gentle stimulation were scored as dead. Lifespan data were analyzed using R software (ver. 4.1.0, “coin” package); Kaplan-Meire survival curves were depicted and *P* values were calculated using the log-rank (Mantel-Cox) test. All lifespan data are described in Extended Data Table 1.

### Triglyceride quantification

Triglyceride (TG) contents were measured using the triglyceride colorimetric assay kit (Abcam, MA, USA) with manufacturer’s protocol. Briefly, frozen worm pellets in liquid nitrogen with 5% Triton X-100. The pellets were sonicated and diluted for protein determination by BCA assay (Pierce, IL, USA). The samples were heated till 80°C and shaken for 5 min then, samples cool-downed to room temperature to solubilize all the triglycerides. TG contents were normalized relative to protein contents and three independently collected worm pellets were assayed for each experimental condition.

### Superoxide dismutase and catalase activity assay

To assess antioxidative enzyme activities, 10-day-old worms grounded in liquid nitrogen. A superoxide dismutase (SOD) and catalase activity levels were measured with the SOD colormetric activity kit (Invitrogen, CA, USA) or Catalase activity colorimetric/fluorometric assay kit (Biovision, CA, USA), respectively. Worm pellet samples were assessed according to manufacturer’s protocol in triplicate. SOD and catalase activity was determined using standard curves followed by normalization to protein concentration using the BCA protein assay (Pierce, IL, USA).

### Stress assays

Synchronized worms treated each condition for 10 days. For the oxidative stress assay, 20 worms were placed on solid NGM plates containing 0.4M paraquat (PESTANAL, Sigma-Aldrich, MO, USA) for 3 hours. Assays repeated for 9 times and the paraquat plates were freshly prepared on the day of the assay. For the thermotolerance assay, 18-28 worms were exposed 35°C for 16 hours and survival worms were counted. Assays performed triplicate and survival rate was estimated.

### Lipofuscin (age pigment) analysis

Nematode were synchronized and treated for 10 days with vehicle, control vector or threonine, *tdh*/F08F3.4 dsRNA from L4 larvae stage (Day 0). Day 10 worms were distributed on a 96well plate (Porvair Sciences black with glass-bottomed imaging plates, #324002). Lipofuscin auto-fluorescence was determined using a fluorescence plate reader (Synergy H1, BioTek, VT, USA; excitation: 390-410, emission: 460-480) with normalized to the stable signal of the worms (excitation: 280-300, emission: 320-340) as blank. For measuring lipofuscin accumulation, worms washed with M9 buffer 3 times and fixed for 10 min in 4% paraformaldehyde, washed again and incubated in 0.1% Triton X-100 for 10 min. After washing, the cover slips were mounted onto glass slides and visualized under a confocal laser scanning microscope (LSM8, Carl Zeiss, Jena, Germany) (with excitation at 350 nm and emission at 420 nm). The fluorescence intensities were calculated densitometrically with ZEN Lite software (Carl Zeiss, Jena, Germany) by measuring the average pixel intensity.

### Movement and body size assay

Day 10 of 10 – 15 worms were transferred to fresh NGM plates and recorded 30 second movement with a microscope system (Olympus SZ61 microscope with Olympus camera eXcope T300, Olympus, Tokyo, Japan). Subsequently, 5 independent movement clips per experimental condition were analyzed with TSView 7 software (ver. 7.1) and average speed was calculated as distance (mm) per second.

For measure body size, Day 10 worms washed with M9 buffer 3 times and fixed for 10 min in 4% paraformaldehyde, washed again and incubated in 0.1% Triton X-100 for 10 min. After washing, the cover slips were mounted onto glass slides and visualized under microscope system (Olympus SZ61 microscope with Olympus camera eXcope T300) and body size was measured with TSView 7 software.

### Fertility assay

Worms were synchronized and single L4 nematode were transferred on single plates applying control or treatment RNAi bacteria or reagent then moved to fresh plates. Progeny of worms were allowed to hatch and counted.

### Pharyngeal pumping rates of *C*.*elegans*

The pharyngeal pumping of 30 worms per each condition were assessed for 1 min using a SZ61 microscope (Olympus, Tokyo, Japan). Pharyngeal contractions were recorded with Olympus camera eXcope T300 at 18-fold optical zoom and videos were played back at 0.5× speed and pharyngeal pumps counted.

### Assay for methylglyoxal (MGO)/Glutathione (GSH) rate in *C. elegans*

For detection of MGO using Methylglyoxal ELISA kit (Abcam, MA, USA), frozen worm pellet was used and sample prepared according to manufacturer’s protocol. Glyoxalase activity was also measured with Glyoxalase I colorimetric assay kit (Abcam, MA, USA). Quantification of GSH in worms was measured using the 5,5’-dithiobis-(2-nitrobenzoic acid) (DTNB) cycling method originally created for cell and tissue samples but optimized for whole worm extracts. Based on protocol ^55^, we detected a 5’-thio-2-nitrobenzoic acid (TNB) chromophore, reaction compound of DTNB and GSH, with maximum absorbance of 412 nm using GSH quantification kit (Dojindo, Kumamoto, Japan).

### DAF-16::GFP / HSF-1::GFP localization assay in *C. elegans*

The *daf-16::gfp* (TJ356) and *hsf-1::gfp* (OG532) strain was used and their fluorescence was visualized using a confocal laser scanning microscope with inverted stand (LSM8, Carl Zeiss, Jena, Germany) (with excitation at 488 nm and emission at 535 nm). For each condition, the nucleus/intermediate/cytoplasm fluorescence or fluorescence intensity of 30 - 40 worms were detected densitometrically with ZEN Lite software (Carl Zeiss, Jena, Germany) with triplicate attempts.

### RNA extraction and quantitative real-time PCR (qRT-PCR) in *C. elegans*

Worms were prepared approximately 500 in three biological replicates per each condition at the desired stage. Total RNA was prepared using RNeasy mini kit (Qiagen, Hilden, Germany) then, RNA concentration and quality was measured with a SPECTROstar Nano (BMG Labtech, GmbH, Germany). cDNAs were prepared using SuperScript III reverse transcript kit (ThermoFisher Scientific, MA, USA). TaqMan RT-PCR technology (7500Fast, Applied Biosystems, CA, USA) was used to determine the expression levels of selected target genes with TaqMan sitespecific primers and probes (Thermo Scientific, MA, USA). The process included a denaturing step performed at 95°C for 10 min followed by 50 cycles of 95°C for 15 s and 60°C for 1 min. The reactions were performed in triplicate and samples were analyzed by ddCT method with normalization to the reference genes *cdc-42* and *Y45F10D*.*4* levels ^56^.

### RNA sequencing (RNA-seq), data analysis, and visualization

RNA was extracted and assigned using Bioanalyzer 2100 (Agilent Technologies, CA, USA) in combination with RNA 6000 nano kit. Matched samples with high RNA integrity number scores were proceed to sequencing. For library preparation an amount of 2 mg of total RNA per sample was processed using Illumina’s TruSeq RNA Sample Prep Kit (Illumina, CA, USA) following the manufacturer’s instruction. Quality and quantity of the libraries was determined using Agilent Bioanalyzer 2100 in combination with FastQC v0.11.7, Sequencing was done on a HiSeq4000 in SR/50 bp/high output mode at the Macrogen Bioinformatics Center (Macrogen, Seoul, Korea).

Libraries were multiplexed in five per lane. Sequencing ends up with 35 Mio reads per sample. Sequence data were extracted in FastQ format using bcl2fastq v1.8.4 (Illumina) and used for mapping approach. FASTQ output files were aligned to the WBcel235 (February 2014) *C. elegans* reference genome using STAR ^57^. These files have been deposited at the Sequence Read Archive (SRA) with the accession number SUB10527691. Samples averaged 75% mapping of sequence reads to the reference genome. We filtered out transcripts with Trimmomatic 0.38 platform ^58^. Differential expression analysis was performed using HISAT2 version 2.1.0, Bowtie2 2.3.4.1, with StringTie version 2.1.3b ^59,60^. Intersection of differentially expressed genes (DEGs) of *tdh/F08F3*.*4* RNAi (padj<0.05 in edgeR, DEseq) was performed with data from indicated published datasets by Venn analysis using the ggVennDiagram package in R software. Human ortholog matching was performed using WormBase, Ensembl, and OrthoList2 ^61^. Gene lists were evaluated for functional classification and statistical overrepresentation with Database for Annotation, Visualization, and Integrated Discovery (DAVID) version 6.8 and Kyoto Encyclopedia of Genes and Genomes (KEGG) pathway database.

### Iron assay

Ion contents were measured using the Iron assay kit (Sigma-Aldrich, MO, USA) with manufacturer’s protocol. Briefly, frozen worm pellets in liquid nitrogen with 5% Triton X-100. The pellets were sonicated with 5 volumes of iron assay buffer and centrifuged at 13,000 g for 10 min at 4°C to remove insoluble material. Released iron by acidic buffer is reacted with a chromagen resulting in a colorimetric (593 nm) product, proportion to the iron present. Ferrous/Ferric iron was determined using standard curves followed by normalization to protein concentration using the BCA protein assay (Pierce, IL, USA). Three biological replicates of the assay were performed.

### Quantification of reactive oxygen species level

Day 10 of 5 plates of worms were washed with M9 buffer and incubated with 100 mM Amplex Red probe (Invitrogen, CA, USA) in Krebs-Ringer Phosphate buffer (145 mM NaCl, 5.7 mM Na_2_PO_4_, 4.86 mM KCl, 0.54 mM CaCl_2_, 1.22 mM MgSO_4_ and 5.5 mM glucose, pH 7.4) containing 0.2 U ml^−1^ of horseradish peroxidase. After occasional mixing in the dark for 3 hours, fluorescence intensity was determined using a fluorescence plate reader (Synergy H1, BioTek, VT, USA; excitation: 571, emission: 585) by normalization using protein input with BCA protein assay (Pierce, IL, USA). Three biological replicates of the measurement were performed.

### Lipid peroxidation assay

For measurement of lipid peroxidation level, we performed a Thiobarbituric acid reactive substances (TBARS) assy kit (Cayman Chemical, MI, USA) to detect malondialdehyde (MDA) level, according to manufacturer’s instructions. Replicate experimental samples were collected and frozen worms homogenized by TissueLyser II (Qiagen, Hilden, Germany) in the 4°C room with condition of 30 Hz for 2 min, then rest on ice for 1 min, repeated 3 times. Lysed worms were centrifuged (13,200 rpm, 15 min, 4°C) to remove debris, and supernatant retained. The protein concentration was determined with BCA protein assay (Pierce, IL, USA) and equivalent 20-25 μg of total protein used for assay. For conditions with acute glutathione depletion, Day 2 animals treated with 20 mM diethyl maleate (DEM) or 5 mM of erastin for 6 hours prior to collection. Three biological replicates of the assay were performed.

## Statistical analysis

All experiments were repeated at least three times with identical or similar results. Data represents biological replicates. Adequate statistical analysis was used for every assays. Data meet the hypothesis of the statistical tests described each experiments. Data are expressed as the mean ± s.d. in all figures unless stated otherwise. R software (ver. 4.1.0) and Excel 2016 (Microsoft, NM, USA) were used for statistical analyses. The *p*-values \**p* < 0.05, \*\**p* < 0.01, \*\*\**p* < 0.001, and ^#^*p* < 0.01 were considered statistically significant.

## Data availability

There is no restriction on data availability. The RNA sequence data used in this study have been deposited in NCBI’s SRA and are accessible through submission number SUB10527692. Source data for all the individual *P* values are provided with this paper.

