## Extended Data Fig. for "L-threonine mediated DAF-16/HSF-1 activation inhibits ferroptosis and increases healthspan"

Juewon Kim, Yunju Jo, Donghyun Cho, Dongryeol Ryu

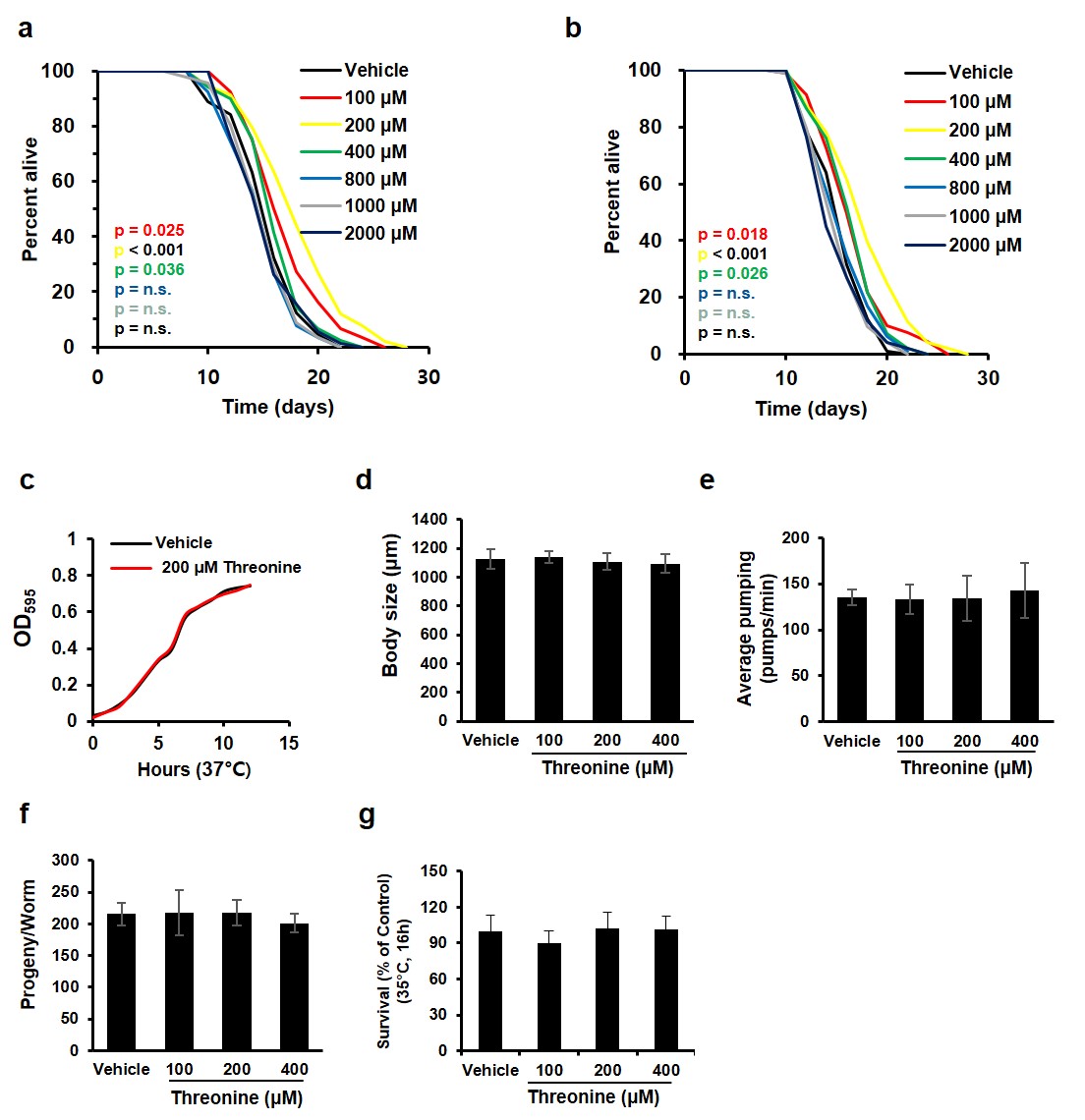

**Extended Data Fig.1 | Threonine prolongs lifespan without diet or fertility changes. a-b**, Survival curves depicted in Fig. 1a with additional replicates (p-values listed, log-rank test). Data of lifespan analysis are displayed in Extended Data Table 1. **c**, Threonine does not alter the growth rate of the OP50 *E. coli*, which is the standard food source for nematodes. **d-g**, Effects of threonine versus the vehicle in animals regarding (**d)** body size (p > 0.5, n = 6), (**e**) average pumping (p > 0.05, Student’s *t*-test, n = 10 worms × 3 assays each), (**f**) progeny (p > 0.05, Student’s *t*-test, three independent measurement), and (**g**) thermotolerance (P > 0.05, Student’s *t*-test, n = 20 worms × 9 measurements each). Error bars represent the mean ± s.d.

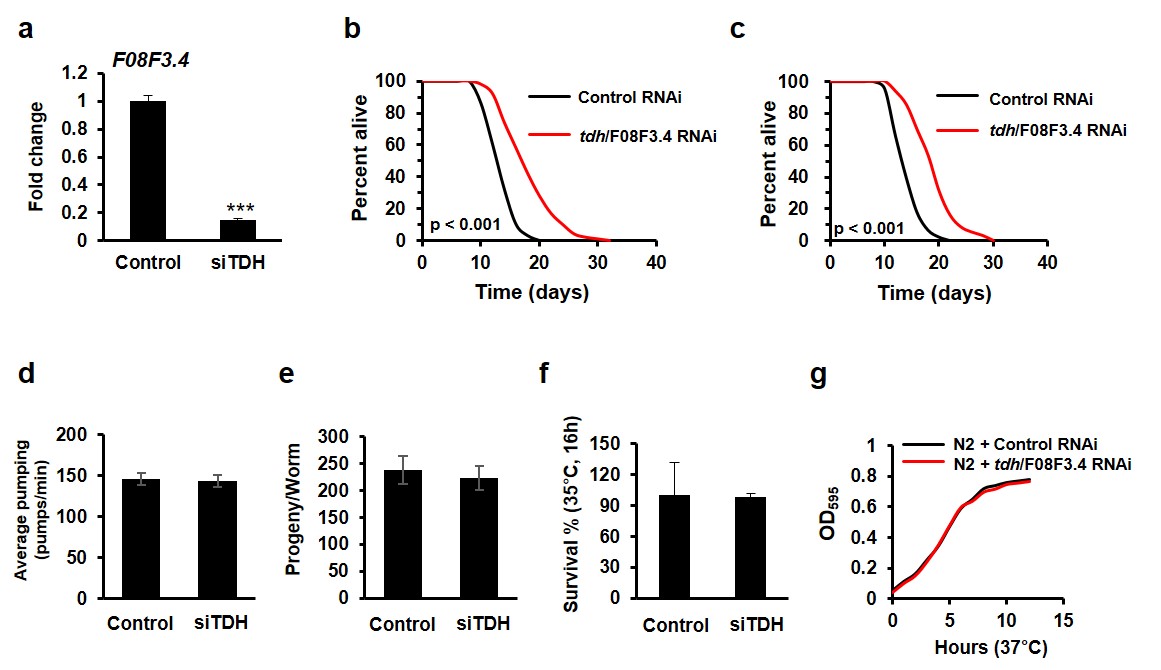

**Extended Data Fig.2 | Downregulation of threonine dehydrogenase extends lifespan without diet or fertility changes.** **a**, The efficiency of *tdh/F08F3.4* knockdown by RNAi was confirmed by quantitative RT-PCR (qRT-PCR) of the mRNA (71% decrease, ***p < 0.001, Student’s *t*-test, three independent measurements). **b-c**, Survival curves depicted in Fig. 2k with additional replicates (p < 0.001, log-rank test). Survival data represented in Extended Data Table 1. **d-g**, Effects of *tdh/F08F3.4* RNAi versus control RNAi-treated animals regarding (**d**) average pumping (p = 0.748, Student’s *t*-test, n = 10 worms × 3 assays each), (**e**) progeny (p = 0.487, Student’s *t*-test, three independent measurement), and (**f**) thermotolerance (p = 0.92, Student’s *t*-test, n = 20 worms × 9 measurements each). **g**, *tdh/F08F3.4* RNAi does not change the growth rate of OP50 *E. coli*. Error bars represent the mean ± s.d.

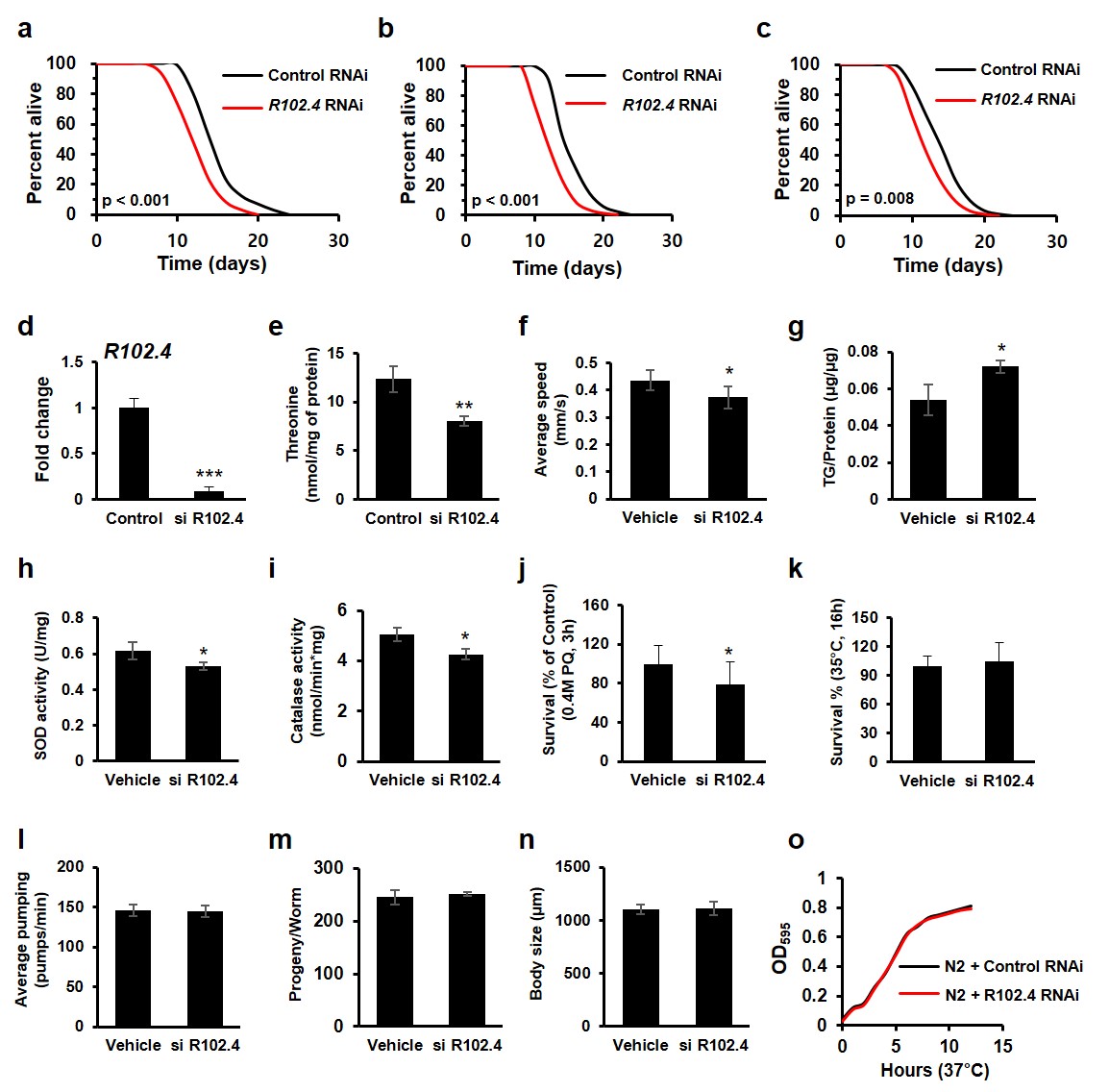

**Extended Data Fig.3 | Downregulation of threonine anabolic enzyme *R102.4* shortens lifespan and reduces healthspan.** **a-c**, Survival curves of R102.4 RNAi versus control RNAi (black) with additional replicates (p < 0.001 or p = 0.008, log-rank test). Lifespan assay data are depicted in Extended Data Table 1. **d**, The efficiency of *R102.4* knockdown by RNAi was confirmed by quantitative RT-PCR (qRT-PCR) of the mRNA (89.9% decrease, ***p < 0.001, Student’s t-test, 3 independent measurement). **e-r**, Effects of *R102.4* RNAi versus control RNAi regarding (**e**) threonine content (**p = 0.006, Student’s *t*-test, n =3 worm pellets), (**f**) average speed (*p = 0.021, Student’s *t*-test, n = 10-15 worms × 3 assays each), (**g**) triglyceride (TG) content (*p = 0.025, Student’s *t*-test, n =3 worm pellets), (**h**) superoxide dismutase (SOD) activity (*p = 0.05, Student’s *t*-test, n = 3 worm pellets), (**i**) catalase activity (**p = 0.014, Student’s *t*-test, n = 3 worm pellets), (**j**) oxidative stress resistance (*p = 0.049, Student’s *t*-test, n = 20 worms × 9 measurements each), (**k**) thermotolerance (p = 0.726, Student’s *t*-test, n = 20 worms × 9 measurements each), (**l**) average pumping (p = 0.852, Student’s *t*-test, n = 10 worms × 3 assays each), (**m**) progeny (p = 0.498, Student’s *t*-test, three independent measurements), and (**n**) body size (p = 0.906, n = 6). (**o**) *R102.4* RNAi does not shift the growth rate of the OP50 *E. coli*. Error bars represent the mean ± s.d.

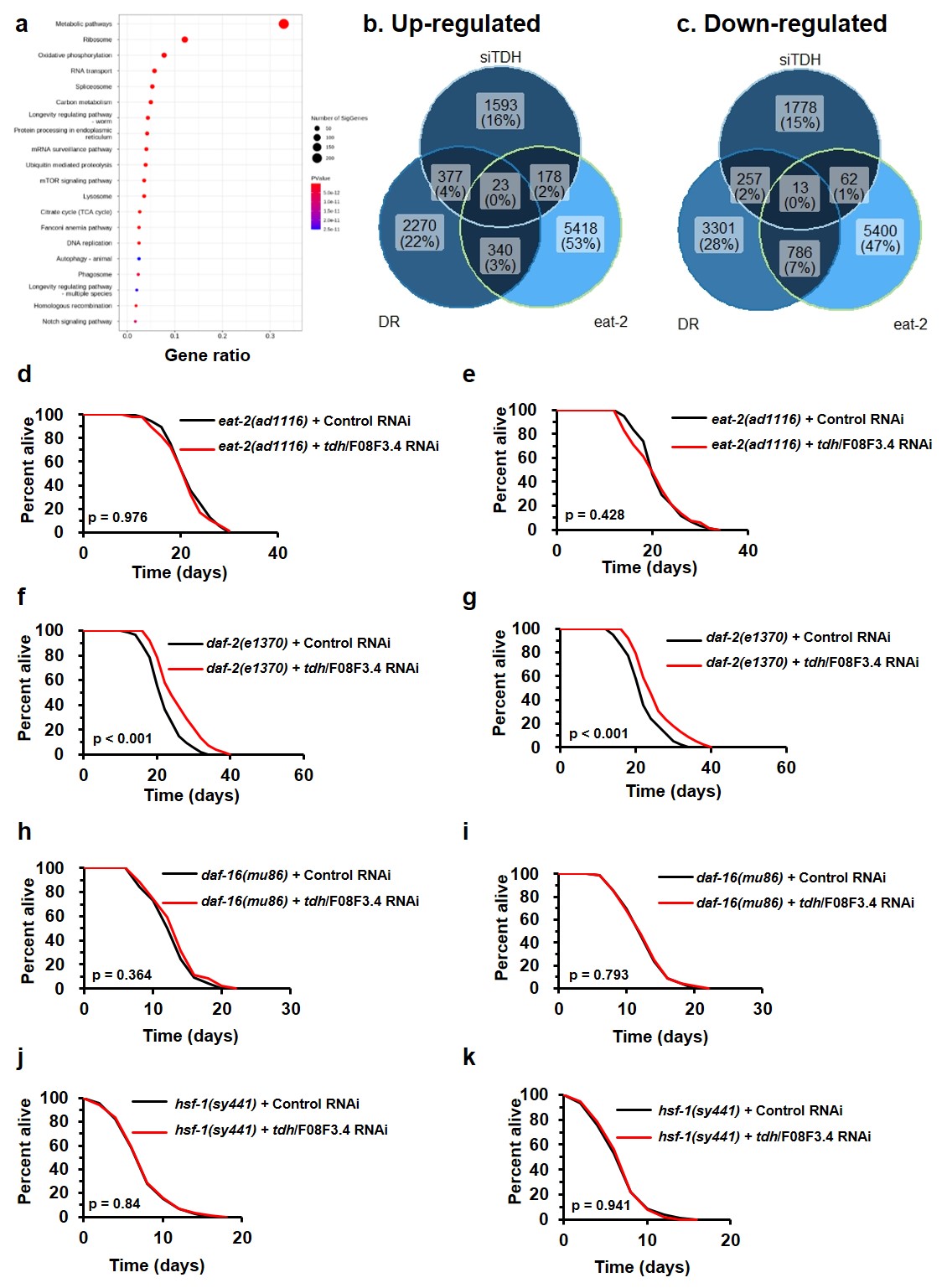

**Extended Data Fig.4 | Downregulation of threonine dehydrogenase alter metabolic process by DAF-16 and HSF-1-mediated mechanism. a**, Functional annotation clustering for the DEGs from the volcano and smear plot of Fig. 3b-c (p < 0.05) determined by gene enrichment analysis using the KEGG database (p < 0.05). **b-c**, Venn analysis of transcripts that are regulated by *tdh/F08F3.4* RNAi (siTDH), dietary restriction (DR), and *eat-2* mutant (eat-2); (**b**) upregulated and (**c**) downregulated genes. **d-e**, Survival curves depicted in Fig. 3d with additional replicates (p-value listed, log-rank test). **f-g**, Survival curves depicted in Fig. 3e with additional replicates (p < 0.001, log-rank test). **h-i**, Survival curves depicted in Fig. 3f with additional replicates (p-value listed, log-rank test). **j-k**, Survival curves depicted in Fig. 3g with additional replicates (p-value listed, log-rank test). Survival data are presented in Extended Data Table 1.

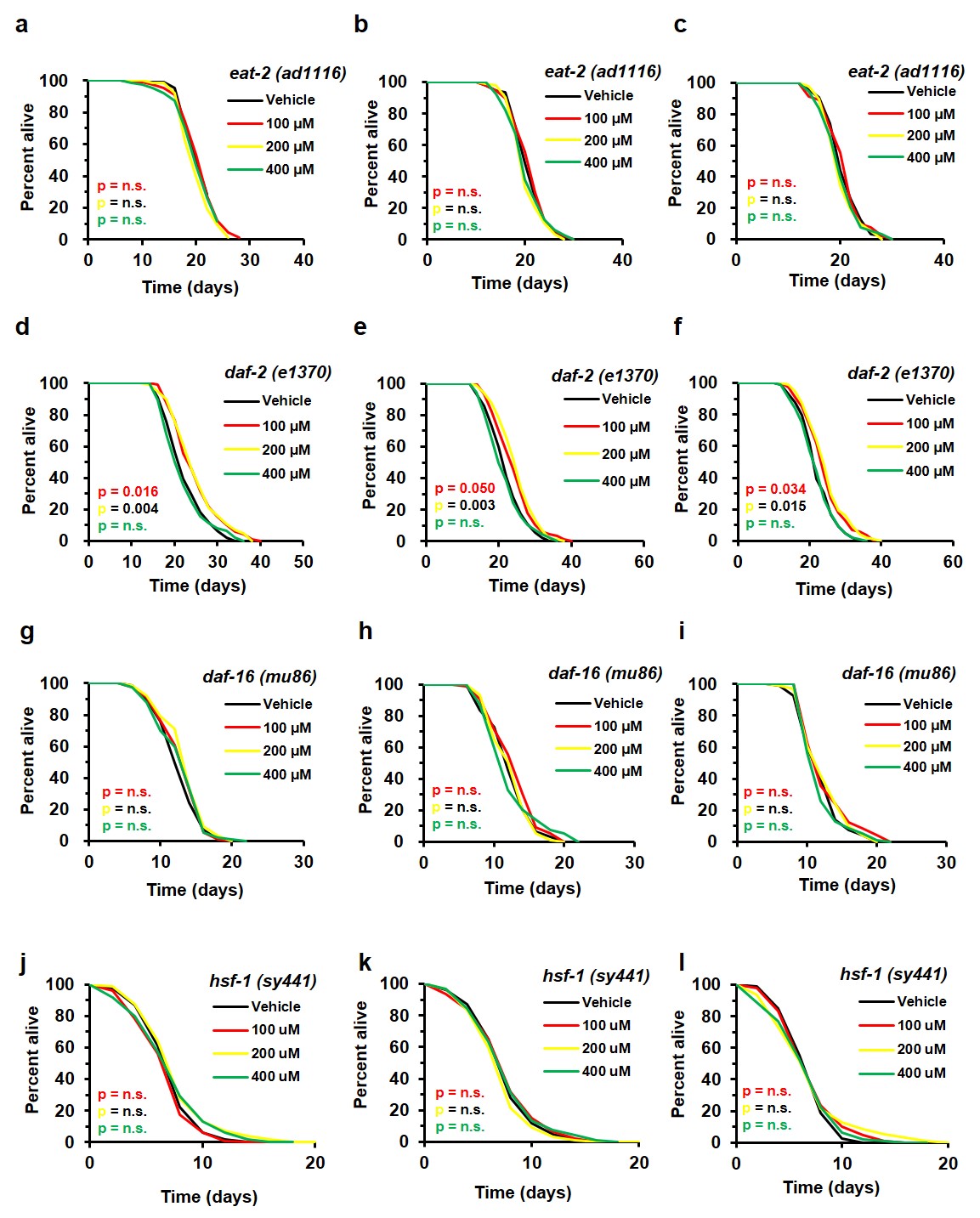

**Extended Data Fig.5 | Threonine supplementation extends lifespan by a DAF-16 and HSF-1-mediated mechanism. a-l**, Effects of L-threonine (100, 200, and 400 μM) versus the vehicle (black) on lifespan of **a-c**, *eat-2*(*ad465*), **d-f**, *daf-2*(*e1370*), **g-i**, *daf-16*(*m26*), and **j-l**, *hsf-1*(*sy441*) with additional replicates; colour coding is assigned to all subsequent panels. P-value and lifespan assay data summarized in Extended Data Table 1.

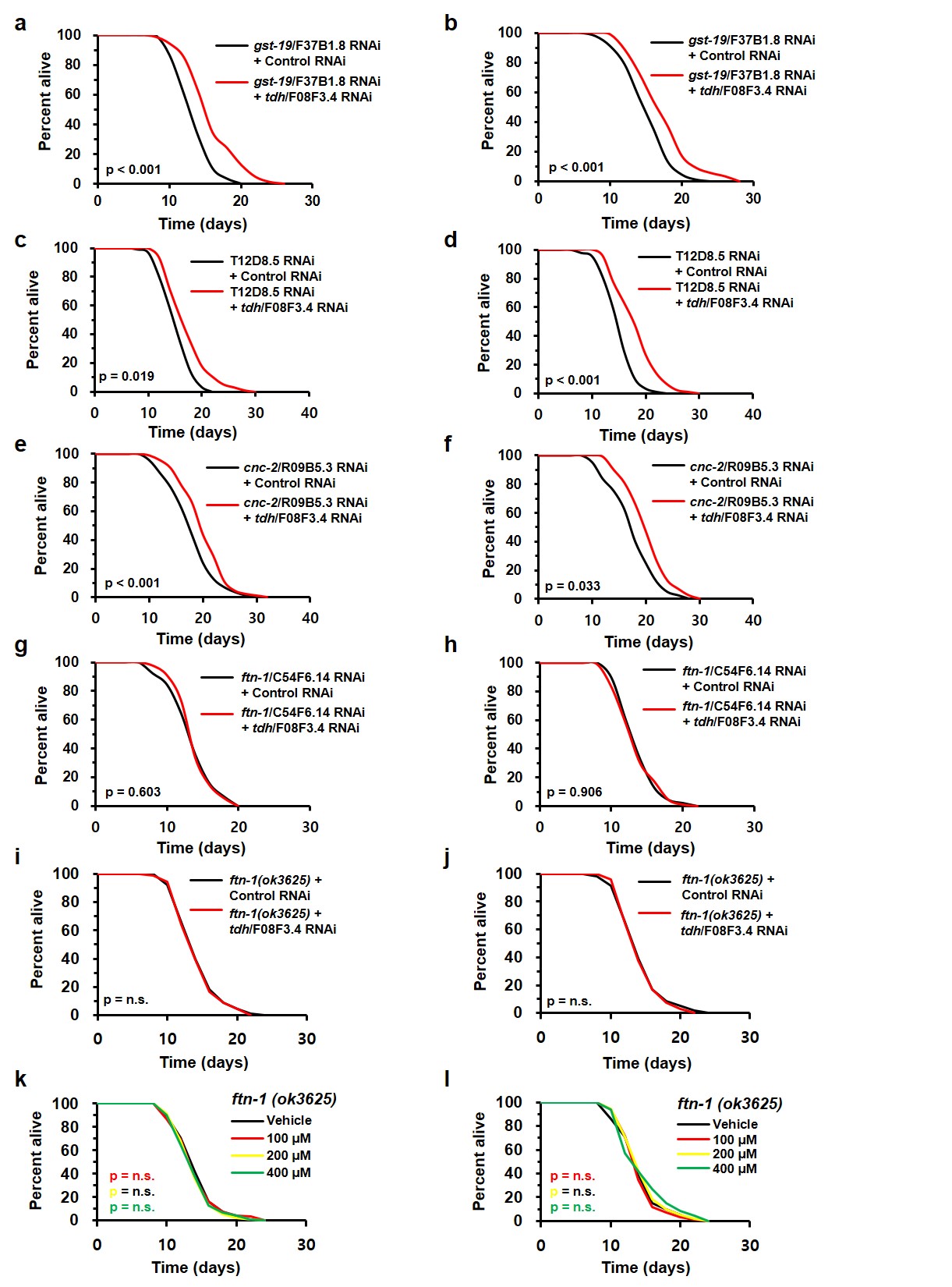

**Extended Data Fig.6 | FTN-1 is necessary for threonine-mediated lifespan extension. a-b**, Survival curves depicted in Fig. 4a with additional replicates. **c-d**, Survival curves depicted in Fig. 4b with additional replicates. **e-f**, Survival curves depicted in Fig. 4c with additional replicates. **g-h**, Survival curves depicted in Fig. 4d with additional replicates. **i-j**, Survival curves depicted in Fig. 4e with additional replicates. **k-l**, Survival curves depicted in Fig. 4f with additional replicates. P-values listed in figure panel and survival data are presented in Extended Data Table 1.

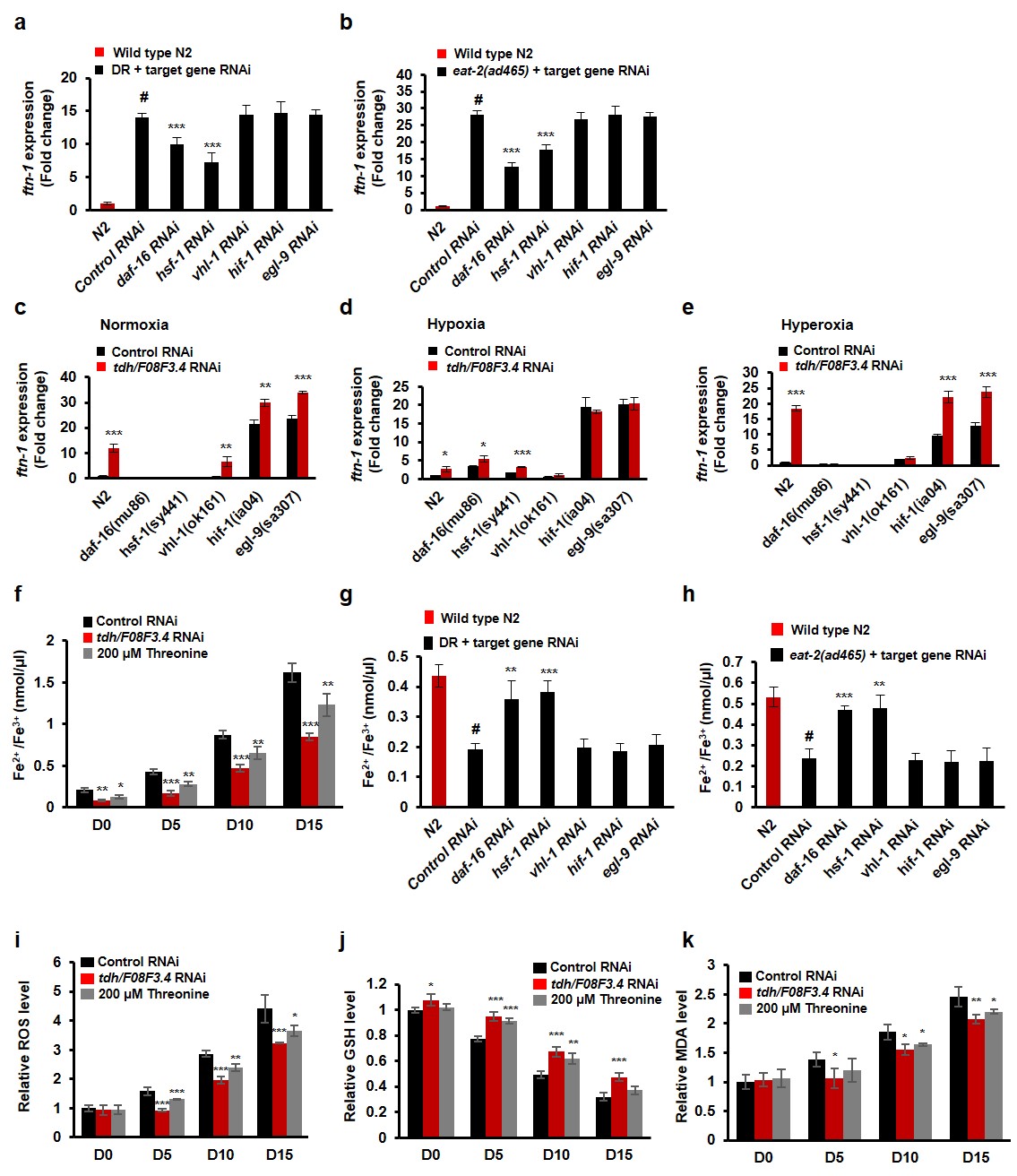

**Extended Data Fig.7 | Increased threonine attenuates ferroptosis by abrogating ROS, lipid peroxidation, and GSH depletion. a-b**, Expression levels of *ftn-1* at **a**, DR, or **b**, *eat-2* mutant with target gene RNAi (#p < 0.001 compared to N2, ***p < 0.001 versus control RNAi, one-way ANOVA, n = 3 worm pellets). **c-e**, Expression of *ftn-1* with *tdh/F08F3.4* RNAi at the various mutants that are related to the regulation of *ftn-1* transcription. Worms were treated under **c**, normoxia (21% O_2_), **d**, hypoxia (2% O_2_), and **e**, hyperoxia (0.4M PQ) for 3 h (*p < 0.05, **p < 0.01, and ***p < 0.001 versus the control RNAi, one-way ANOVA, n =3 worm pellets). **f**, Quantification of Fe^2+^ /Fe^3+^ iron contents of nematode at intervals across lifespan (*p < 0.05, **p < 0.01, and ***p < 0.001 versus the vehicle group, one-way ANOVA, n = 3 worm pellets). **g-h**, Fe^2+^ /Fe^3+^ iron content levels of (**g**) DR or (**h**) *eat-2* mutants with the target gene RNAi were measured (#p < 0.001 compared to N2, **p < 0.01 and ***p < 0.001 versus control RNAi, one-way ANOVA, n =3 worm pellets). **i**, Relative Amplex Red fluorescence in supernatant of worms (*p < 0.05, **p < 0.01, and ***p < 0.001 versus control group, one-way ANOVA, n = 3 worm pellets). **j**, Total glutathione (GSH) level was normalized to the GSH level in worms not exposed to *tdh/F08F3.4* RNAi or threonine (*p < 0.05, **p < 0.01, and ***p < 0.001 versus control, one-way ANOVA, n = 3 worm pellets). **k**, Levels of the lipid peroxidation end product, malondialdehyde (MDA), were measured and normalized against the mean of untreated worms for independent samples (*p < 0.05 and **p < 0.05 versus control, one-way ANOVA, n = 3 worm pellets).

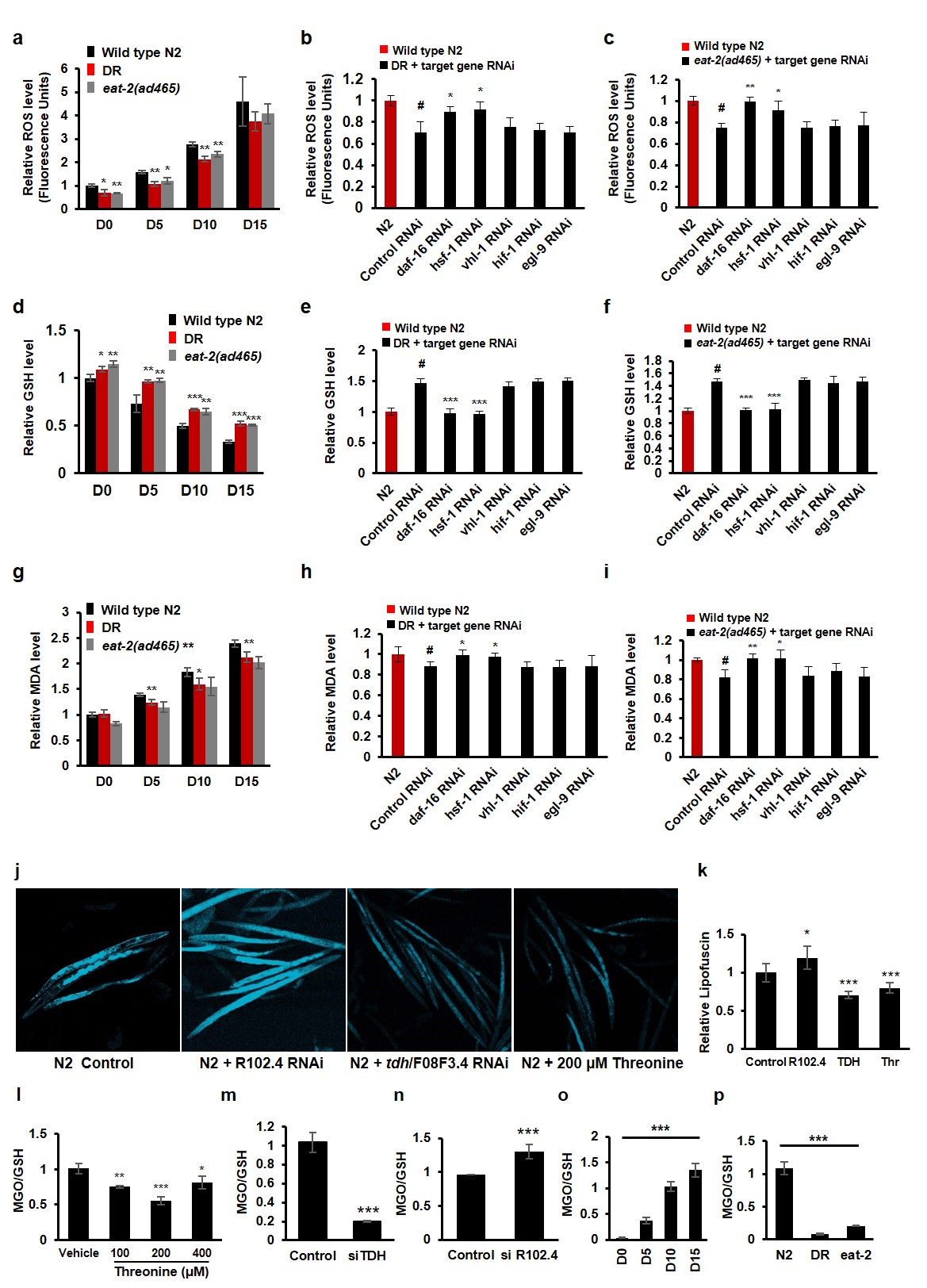

**Extended Data Fig.8 | DR represented decreased ferroptosis in the manner of DAF-16 and HSF-1 and increasing threonine lowered age-associated factors. a**, Relative Amplex Red fluorescence in supernatant of worm strains at intervals across lifespan (*p < 0.05 and **p < 0.01 versus the N2 group, one-way ANOVA, n = 3 worm pellets). **b-c**, Relative ROS level of (**b**) DR and (**c**) *eat-2* mutant with target gene RNAi (#p < 0.001 compared to N2, *p < 0.05 and **p < 0.01 versus the control RNAi, one-way ANOVA, n = 3 worm pellets). **d**, Relative GSH levels of worm strains at intervals across lifespan (*p < 0.05, **p < 0.01, and ***p < 0.001 versus N2, one-way ANOVA, n = 3 worm pellets). **e-f**, Relative GSH level of (**e**) DR and (**f**) *eat-2* mutant with target gene RNAi (#p < 0.001 compared to N2, ***p < 0.001 versus the control RNAi, one-way ANOVA, n = 3 worm pellets). **g**, Relative MDA levels of worm strains at intervals across lifespan (*p < 0.05 and **p < 0.01 versus N2, one-way ANOVA, n = 3 worm pellets). **h-i**, Relative MDA in (**h**) DR and (**i**) *eat-2* mutant with target gene RNAi (#p < 0.001 compared to N2, *p < 0.05 and **p < 0.01 versus control the RNAi, one-way ANOVA, n = 3 worm pellets). **j**, The auto-fluorescence of lipofuscin and **k**, Relative lipofuscin levels with *R102.4* RNAi, *tdh/F08F3.4* RNAi, and 200 μM threonine treatment (*p < 0.05 and ***p < 0.001 versus the control, one-way ANOVA, n = 3 worm pellet). **l-p**, MGO/GSH levels under (**l**) threonine treatment (*p < 0.05, **p < 0.01, and ***p < 0.001 versus the vehicle, one-way ANOVA, n = 3 worm pellet), (**m**) *tdh/F08F3.4* RNAi, (**n**) *R102.4* RNAi (***p < 0.001 versus control RNAi, Student’s t-test, n =3 worm pellets), (**o**) interval time points (***p < 0.001 versus Day 0, one-way ANOVA, n = 3 worm pellet), and (**p**) N2, DR, and *eat-2* mutant (***p < 0.001 versus N2, one-way ANOVA, n = 3 worm pellet). Error bars represent the mean ± s.d.

**Extended Data Table 1 | Summary of lifespan data**

| Strain | Treatment | Mean lifespan (days) ± SEM | | % difference | P-value | n (animals) | Figure |
| --- | --- | --- | --- | --- | --- | --- | --- |
| N2 |  | Vehicle | 15.7 ± 0.3 |  |  | 90 | Fig. 1c |
|  | Threonine (μM) | 100 | 17.2 ± 0.4 | 9.5 | 0.01318723 | 90 |  |
|  |  | 200 | 18.5 ± 0.4 | 18 | 9.818E-06 | 87 |  |
|  |  | 400 | 16.5 ± 0.3 | 5.4 | 0.02632925 | 89 |  |
|  |  | 800 | 15.1 ± 0.3 | -3.5 | 0.29197009 | 88 |  |
|  |  | 1000 | 15.5 ± 0.3 | -1.3 | 0.47151619 | 90 |  |
|  |  | 2000 | 15.4 ± 0.3 | -1.9 | 0.39110771 | 83 |  |
| N2 |  | Vehicle | 15.7 ± 0.3 |  |  | 90 | Extended data Fig. 1a |
|  | Threonine (μM) | 100 | 17.4 ± 0.4 | 10.8 | 0.025 | 92 |  |
|  |  | 200 | 18.4 ± 0.4 | 17.3 | 4.0351E-05 | 92 |  |
|  |  | 400 | 16.5 ± 0.3 | 5 | 0.03624563 | 89 |  |
|  |  | 800 | 15.2 ± 0.3 | -3.1 | 0.24118061 | 93 |  |
|  |  | 1000 | 15.4 ± 0.3 | -1.9 | 0.24921714 | 91 |  |
|  |  | 2000 | 15.6 ± 0.3 | -0.8 | 0.29940049 | 91 |  |
| N2 |  | Vehicle | 15.8 ± 0.3 |  |  | 89 | Extended data Fig. 1b |
|  | Threonine (μM) | 100 | 17.1 ± 0.4 | 8.7 | 0.01828145 | 91 |  |
|  |  | 200 | 18.2 ± 0.4 | 15.7 | 2.8716E-05 | 96 |  |
|  |  | 400 | 16.9 ± 0.3 | 7.4 | 0.02617666 | 96 |  |
|  |  | 800 | 15.9 ± 0.3 | 0.9 | 0.60101601 | 95 |  |
|  |  | 1000 | 15.5 ± 0.3 | -1.8 | 0.695817 | 93 |  |
|  |  | 2000 | 15.3 ± 0.3 | -2.7 | 0.56453947 | 93 |  |
| N2 | Control RNAi |  | 14.6 ± 0.3 |  |  | 92 | 2k |
|  | TDH RNAi |  | 19.5 ± 0.4 | 33.4 | 4.1701E-14 | 96 |  |
| N2 | Control RNAi |  | 15.8 ± 0.3 |  |  | 88 | Extended data Fig. 2b |
|  | TDH RNAi |  | 21.5 ± 0.5 | 36.1 | 1.5979E-09 | 90 |  |
| N2 | Control RNAi |  | 15.2 ± 0.3 |  |  | 91 | Extended data Fig. 2c |
|  | TDH RNAi |  | 20.3 ± 0.4 | 33.6 | 3.5476E-11 | 94 |  |
| N2 | Control RNAi |  | 15.6 ± 0.3 |  |  | 96 | Extended data Fig. 3a |
|  | R102.4 RNAi |  | 13.0 ± 0.2 | -16.8 | 0.00015631 | 94 |  |
| N2 | Control RNAi |  | 15.7 ± 0.3 |  |  | 90 | Extended data Fig. 3b |
|  | R102.4 RNAi |  | 13.1 ± 0.2 | -16.5 | 3.6303E-06 | 95 |  |
| N2 | Control RNAi |  | 14.7 ± 0.3 |  |  | 90 | Extended data Fig. 3c |
|  | R102.4 RNAi |  | 12.7 ± 0.3 | -13.6 | 0.00875291 | 95 |  |
| *eat-2(ad1116)* | Control RNAi |  | 21.7 ± 0.4 |  |  | 102 | 3d |
|  | TDH RNAi |  | 21.6 ± 0.4 | -0.4 | 0.785831 | 97 |  |
| *eat-2(ad1116)* | Control RNAi |  | 21.8 ± 0.4 |  |  | 94 | Extended data Fig. 4d |
|  | TDH RNAi |  | 21.2 ± 0.5 | -2.6 | 0.97644574 | 94 |  |
| *eat-2(ad1116)* | Control RNAi |  | 21.4 ± 0.5 |  |  | 93 | Extended data Fig. 4e |
|  | TDH RNAi |  | 20.9 ± 0.5 | -2.3 | 0.42830439 | 96 |  |
| *eat-2(ad1116)* |  | Vehicle | 21.3 ± 0.4 |  |  | 91 | Extended data Fig. 5a |
|  | Threonine (μM) | 100 | 21.1 ± 0.4 | -0.7 | 0.82883163 | 91 |  |
|  |  | 200 | 20.4 ± 0.3 | -3.8 | 0.19526359 | 88 |  |
|  |  | 400 | 20.7 ± 0.4 | -2.9 | 0.91731597 | 88 |  |
| *eat-2(ad1116)* |  | Vehicle | 21.0 ± 0.3 |  |  | 98 | Extended data Fig. 5b |
|  | Threonine (μM) | 100 | 21.2 ± 0.4 | 0.8 | 0.47588931 | 91 |  |
|  |  | 200 | 20.4 ± 0.3 | -2.8 | 0.3479877 | 94 |  |
|  |  | 400 | 20.5 ± 0.4 | -2.3 | 0.69626906 | 97 |  |
| *eat-2(ad1116)* |  | Vehicle | 21.0 ± 0.4 |  |  | 96 | Extended data Fig. 5c |
|  | Threonine (μM) | 100 | 21.0 ± 0.4 | 0.3 | 0.61436981 | 90 |  |
|  |  | 200 | 20.4 ± 0.3 | -2.7 | 0.31910299 | 94 |  |
|  |  | 400 | 21.0 ± 0.7 | -2.6 | 0.78454987 | 93 |  |
| *daf-2(e1370)* | Control RNAi |  | 22.5 ± 0.5 |  |  | 93 | 3e |
|  | TDH RNAi |  | 25.9 ± 0.6 | 15.1 | 0.00024409 | 97 |  |
| *daf-2(e1370)* | Control RNAi |  | 22.2 ± 0.5 |  |  | 96 | Extended data Fig. 4f |
|  | TDH RNAi |  | 25.8 ± 0.6 | 16.2 | 0.00021323 | 96 |  |
| *daf-2(e1370)* | Control RNAi |  | 22.3 ± 0.5 |  |  | 95 | Extended data Fig. 4g |
|  | TDH RNAi |  | 25.5 ± 0.6 | 14.6 | 0.00222086 | 102 |  |
| *daf-2(e1370)* |  |  | 22.6 ± 0.5 |  |  | 92 | Extended data Fig. 5d |
|  | Threonine (μM) | 100 | 25.1 ± 0.5 | 10.9 | 0.01662628 | 97 |  |
|  |  | 200 | 25.3 ± 0.6 | 11.7 | 0.00382294 | 100 |  |
|  |  | 400 | 22.2 ± 0.5 | -1.8 | 0.87212671 | 96 |  |
| *daf-2(e1370)* |  |  | 22.3 ± 0.5 |  |  | 92 | Extended data Fig. 5e |
|  | Threonine (μM) | 100 | 24.5 ± 0.5 | 9.5 | 0.05007683 | 94 |  |
|  |  | 200 | 25.2 ± 0.5 | 12.6 | 0.00273856 | 97 |  |
|  |  | 400 | 21.9 ± 0.5 | -2.1 | 0.70293411 | 94 |  |
| *daf-2(e1370)* |  |  | 22.7 ± 0.5 |  |  | 89 | Extended data Fig. 5f |
|  | Threonine (μM) | 100 | 24.5 ± 0.6 | 8 | 0.03400012 | 86 |  |
|  |  | 200 | 25.0 ± 0.5 | 10.4 | 0.01544843 | 92 |  |
|  |  | 400 | 21.9 ± 0.5 | -3.2 | 0.83582142 | 88 |  |
| *daf-16(mu86)* | Control RNAi |  | 13.0 ± 0.3 |  |  | 102 | 3f |
|  | TDH RNAi |  | 13.5 ± 0.3 | 0.9 | 0.62725723 | 99 |  |
| *daf-16(mu86)* | Control RNAi |  | 12.9 ± 0.3 |  |  | 97 | Extended data Fig. 4h |
|  | TDH RNAi |  | 13.5 ± 0.3 | 4.5 | 0.36382365 | 95 |  |
| *daf-16(mu86)* | Control RNAi |  | 12.8 ± 0.3 |  |  | 93 | Extended data Fig. 4i |
|  | TDH RNAi |  | 12.8 ± 0.3 | 0.6 | 0.79278221 | 92 |  |
| *daf-16(mu86)* |  |  | 12.9 ± 0.3 |  |  | 87 | Extended data Fig. 5g |
|  | Threonine (μM) | 100 | 13.3 ± 0.3 | 2.9 | 0.48525786 | 89 |  |
|  |  | 200 | 13.7 ± 0.3 | 6.4 | 0.16755809 | 87 |  |
|  |  | 400 | 13.2 ± 0.3 | 1.8 | 0.34965062 | 90 |  |
| *daf-16(mu86)* |  |  | 12.7 ± 0.3 |  |  | 92 | Extended data Fig. 5h |
|  | Threonine (μM) | 100 | 13.2 ± 0.3 | 4.6 | 0.50873332 | 97 |  |
|  |  | 200 | 12.7 ± 0.3 | 0.5 | 0.50153425 | 93 |  |
|  |  | 400 | 12.5 ± 0.4 |  | 0.85684873 | 94 |  |
| *daf-16(mu86)* |  |  | 12.3 ± 0.3 |  |  | 93 | Extended data Fig. 5i |
|  | Threonine (μM) | 100 | 13.0 ± 0.3 | 5 | 0.32790939 | 95 |  |
|  |  | 200 | 12.7 ± 0.3 | 3.2 | 0.4104227 | 95 |  |
|  |  | 400 | 12.2 ± 0.3 | -1 | 0.65040023 | 92 |  |
| *hsf-1 (sy441)* | Control RNAi |  | 8.0 ± 0.3 |  |  | 80 | 3g |
|  | TDH RNAi |  | 8.2 ± 0.3 | 2.7 | 0.49201391 | 83 |  |
| *hsf-1 (sy441)* | Control RNAi |  | 7.8 ± 0.3 |  |  | 86 | Extended data Fig. 4j |
|  | TDH RNAi |  | 7.8 ± 0.2 | 1 | 0.83963669 | 90 |  |
| *hsf-1 (sy441)* | Control RNAi |  | 7.2 ± 0.2 |  |  | 91 | Extended data Fig. 4k |
|  | TDH RNAi |  | 7.2 ± 0.3 | 1 | 0.94061687 | 89 |  |
| *hsf-1 (sy441)* |  |  | 7.5 ± 0.2 |  |  | 98 | Extended data Fig. 5j |
| *hsf-1 (sy441)* | Threonine (μM) | 100 | 7.1 ± 0.2 | -5.7 | 0.54254055 | 98 |  |
| *hsf-1 (sy441)* |  | 200 | 8.1 ± 0.3 | 7.6 | 0.28327525 | 98 |  |
| *hsf-1 (sy441)* |  | 400 | 7.6 ± 0.2 | 0.9 | 0.28874204 | 99 |  |
| *hsf-1 (sy441)* |  |  | 7.9 ± 0.3 |  |  | 86 | Extended data Fig. 5k |
| *hsf-1 (sy441)* | Threonine (μM) | 100 | 7.5 ± 0.3 | 0.9 | 0.58359758 | 95 |  |
| *hsf-1 (sy441)* |  | 200 | 7.5 ± 0.3 | -5.1 | 0.54965683 | 89 |  |
| *hsf-1 (sy441)* |  | 400 | 8.0 ± 0.4 | 2 | 0.70293555 | 96 |  |
| *hsf-1 (sy441)* |  |  | 7.2 ± 0.2 |  |  | 81 | Extended data Fig. 5l |
| *hsf-1 (sy441)* | Threonine (μM) | 100 | 7.4 ± 0.2 | 3.3 | 0.4769379 | 90 |  |
| *hsf-1 (sy441)* |  | 200 | 7.4 ± 0.2 | 2.9 | 0.38189565 | 93 |  |
| *hsf-1 (sy441)* |  | 400 | 7.0 ± 0.2 | -3.1 | 0.42616027 | 99 |  |
| N2 | Control RNAi |  | 14.8 ± 0.3 |  |  | 88 | 4a |
|  | gst-19 RNAi |  | 16.3 ± 0.4 | 18.1 | 0.00011133 | 87 |  |
| N2 | Control RNAi |  | 15.3 ± 0.3 |  |  | 87 | Extended data Fig. 6a |
|  | gst-19 RNAi |  | 18.5 ± 0.5 | 21.1 | 1.5979E-09 | 90 |  |
| N2 | Control RNAi |  | 15.6 ± 0.3 |  |  | 90 | Extended data Fig. 6b |
|  | gst-19 RNAi |  | 17.7 ± 0.4 | 13.8 | 0.01044407 | 89 |  |
| N2 | Control RNAi |  | 15.7 ± 0.3 |  |  | 91 | 4b |
|  | T12D8.5 RNAi |  | 20.5 ± 0.5 | 13.2 | 0.01854408 | 95 |  |
| N2 | Control RNAi |  | 15.5 ± 0.2 |  |  | 91 | Extended data Fig. 6c |
|  | T12D8.5 RNAi |  | 19.8 ± 0.5 | 21.2 | 6.7349E-09 | 92 |  |
| N2 | Control RNAi |  | 15.3 ± 0.3 |  |  | 85 | Extended data Fig. 6d |
|  | T12D8.5 RNAi |  | 20.6 ± 0.4 | 15.6 | 0.00980001 | 88 |  |
| N2 | Control RNAi |  | 18.4 ± 0.4 |  |  | 85 | 4c |
|  | cnc-2 RNAi |  | 20.5 ± 0.5 | 11.1 | 0.02717264 | 87 |  |
| N2 | Control RNAi |  | 18.2 ± 0.4 |  |  | 89 | Extended data Fig. 6e |
|  | cnc-2 RNAi |  | 20.4 ± 0.4 | 12.1 | 0.00148891 | 91 |  |
| N2 | Control RNAi |  | 18.0 ± 0.4 |  |  | 85 | Extended data Fig. 6f |
|  | cnc-2 RNAi |  | 20.6 ± 0.4 | 14.4 | 0.03255776 | 88 |  |
| N2 | Control RNAi |  | 14.0 ± 0.2 |  |  | 92 | 4d |
|  | ftn-1 RNAi |  | 14.0 ± 0.3 | -0.2 | 0.75088368 | 89 |  |
| N2 | Control RNAi |  | 14.0 ± 0.3 |  |  | 90 | Extended data Fig. 6g |
|  | ftn-1 RNAi |  | 14.3 ± 0.3 | 2.4 | 0.60254214 | 90 |  |
| N2 | Control RNAi |  | 14.0 ± 0.2 |  |  | 95 | Extended data Fig. 6h |
|  | ftn-1 RNAi |  | 13.8 ± 0.2 | -1.4 | 0.90572168 | 94 |  |
| *ftn-1 (ok3625)* | Control RNAi |  | 14.5 ± 0.3 |  |  | 100 | 4e |
|  | TDH RNAi |  | 14.5 ± 0.3 | 0.3 | 0.82261213 | 102 |  |
| *ftn-1 (ok3625)* | Control RNAi |  | 14.6 ± 0.3 |  |  | 93 | Extended data Fig. 6i |
|  | TDH RNAi |  | 14.5 ± 0.3 | -0.6 | 0.9317674 | 92 |  |
| *ftn-1 (ok3625)* | Control RNAi |  | 14.6 ± 0.3 |  |  | 96 | Extended data Fig. 6j |
|  | TDH RNAi |  | 14.5 ± 0.3 | -0.1 | 0.68241871 | 95 |  |
| *ftn-1 (ok3625)* |  |  | 14.5 ± 0.3 |  |  | 89 | 4f |
|  | Threonine (μM) | 100 | 14.5 ± 0.3 | -0.38 | 0.83269299 | 91 |  |
|  |  | 200 | 14.4 ± 0.3 | -1.66 | 0.8348663 | 95 |  |
|  |  | 400 | 14.6 ± 0.4 | 0.5 | 0.89492254 | 92 |  |
| *ftn-1 (ok3625)* |  |  | 14.6 ± 0.3 |  |  | 93 | Extended data Fig. 6k |
|  | Threonine (μM) | 100 | 14.5 ± 0.3 | -0.6 | 0.86549436 | 92 |  |
|  |  | 200 | 14.3 ± 0.3 | -1.6 | 0.50980404 | 92 |  |
|  |  | 400 | 14.4 ± 0.3 | -1.4 | 0.68673723 | 92 |  |
| *ftn-1 (ok3625)* |  |  | 14.5 ± 0.3 |  |  | 92 | Extended data Fig. 6l |
|  | Threonine (μM) | 100 | 14.5 ± 0.3 | -0.4 | 0.38539193 | 95 |  |
|  |  | 200 | 14.9 ± 0.3 | 2.4 | 0.92239689 | 92 |  |
|  |  | 400 | 14.9 ± 0.4 | 2.9 | 0.74442601 | 93 |  |

**Extended Data Table 2 | Sequences of qPCR primers**

| Primer | Sequence_Fwd | Sequence_Rev |
| --- | --- | --- |
| *tdh* | TCGCGTTAACGCTAGCATGGATCTC | GTAACATCAGAGATTTTGAGACAC |
| *R102.4* | GGCGAGGAGATAATCGTCGG | GTGACAATCGGGTATACTCGTCA |
| *gst-19* | TCGCGTTAACGCTAGCATGGATCTC | GTAACATCAGAGATTTTGAGACAC |
| *T12D8.5* | TCGCGTTAACGCTAGCATGGATCTC | GTAACATCAGAGATTTTGAGACAC |
| *cnc-2* | TCGCGTTAACGCTAGCATGGATCTC | GTAACATCAGAGATTTTGAGACAC |
| *ftn-1* | TCGCGTTAACGCTAGCATGGATCTC | GTAACATCAGAGATTTTGAGACAC |
| *daf-16* | GCGAATCGGTTCCAGCAATTCCAA | ATCCACGGACACTGTTCAACTCGT |
| *hsf-1* | GGAAAGTGGTCCACATCGAG | TTCACTCTCCCGCAGGATGG |
| *hif-1* | CAGTGATTCTTCAATTCTTTACGTC | GGATTAACACAGACAGATTTAACAG |
| *egl-9* | GCCGACTTTCAATCCACTTC | AATGATCGGAGATCGACTGG |
| *actin* | GAGAGGGAAATCGTGCGTGAC | CATCTGCTGGAAGGTGGACA |
| *cdc-42* | CTGCTGGACAGGAAGATTACG | CTCGGACATTCTCGAATGAAG |
| *Y45F10D.4* | GTCGCTTCAAATCAGTTCAG | GTTCTTGTCAAGTGATCCGACA |
